# Flying in the Face of Adversity: A Drosophila-based Virtual CURE Provides Semester-long Authentic Research Opportunity to the Flipped Classroom

**DOI:** 10.1101/2021.06.28.450232

**Authors:** Edward A. Waddell, Dara Ruiz-Whalen, Alana M. O’Reilly, Nathan T. Fried

## Abstract

A call for the integration of research experiences into all biology curricula has been a major goal for educational reform efforts nationally. Course-Based Undergraduate Research Experiences (CUREs) have been the predominant method of accomplishing this, but their associated costs and complex design can limit their wide adoption. In 2020, the COVID-19 pandemic forced programs to identify unique ways to still provide authentic research experiences while students were virtual. We report here a full guide for the successful implementation of a semester-long virtual CURE that uses *Drosophila* behavioral assays to explore the connection between pain and addiction with the use of a “lab-in-a-box” sent home to students. Individual components were piloted across three semesters and launched as a 100-level introductory course with 19 students. We found that this course increased science identity and successfully improved key research competencies as per the Undergraduate Research Student Self-Assessment (URSSA) survey. This course is ideal for flipped classrooms ranging from introductory biology to upper-level neuroscience courses and can be integrated directly into the lecture period without the need for building a new course. Given the low cost, recent comfort with virtual learning environments, and the current proliferation of flipped biology classrooms following the 2020 pandemic, this curriculum could serve as an ideal project-based active-learning tool for equitably increasing access to authentic research experiences.

## Introduction

For students to develop critical thinking skills as learners, scientists, and citizens, they must participate in activities that apply both practical research techniques and the scientific process in real-world contexts (1, 2). Undergraduates can accomplish this through mentored research experiences which are a major goal of educational reform efforts. These are integral to biology education as they provide a high academic challenge, active collaborative and experiential learning, enriching educational experience, intense student-faculty interaction, and a supportive campus environment (3). These activities, however, are not widely available to all students due to high student-faculty ratios and challenges integrating authentic research experiences into the biology curricula. Given the growing comfort-level with virtual learning environments post-COVID and the subsequent wide movement toward pre-recorded lectures, a unique opportunity is presented to integrate a project-based authentic research experience directly into the classroom’s weekly three-hour lecture period in either an in-person or virtual setting. By strategically focusing on behavioral assays as experimental outputs, costs can also be dramatically reduced.

Considerable research has demonstrated the benefits of undergraduate research experiences on student learning, including the development of domain expertise, acquisition of team-based skills, increased understanding and respect for the research process, acquisition of problem-solving skills, practice and refinement of communication skills, and increased self-confidence, personal growth, independence, and tolerance (4–10). Comparable benefits are seen across race and gender and across various institutional types including research universities, Master’s-level institutions, and teaching colleges (11). Furthermore, these experiences are hypothesized to especially benefit women and underrepresented students due to the fostering of mentor-mentee and peer-peer relationships (12–14).

Despite these recommendations, however, the practicalities of expanding this to an entire undergraduate population is extremely daunting (15, 16). Historically, the incorporation of research experiences into the undergraduate curriculum has been in the form of mentored one-on-one research apprenticeships in faculty research laboratories (5, 11). However, the high student-faculty ratio often prevents all students from participating, often at the cost of diversity and equity in training (17, 18). A solution to this problem is to integrate authentic research experiences into a course-based setting.

Course-based undergraduate research experiences (CUREs) are a scalable solution for providing authentic research experiences to undergraduate students while still fulfilling the benchmarks set by the National Research Council (8, 19). CUREs offer the capacity to involve a greater number of students with diverse backgrounds in research for credit towards their degrees, not just self-selected students who seek out individual research opportunities (20). This is especially the case for students of low socioeconomic backgrounds who often are unable to participate in extracurricular research activities due to employment. Additionally, CUREs can be integrated into introductory-level courses to engage first- and second-year students in research much earlier and thus have the potential to exert a larger influence on students’ academic and career choices (21).

Despite the numerous implementations and successes of CUREs, they are still widely underused. Since CUREs are generally offered in a teaching laboratory versus an established research laboratory, a dedicated laboratory space to create a scalable, authentic research experience may not be feasible for some institutions. Further, CUREs are often stand-alone electives which can disincentive programs adopting them because they can be perceived as too cumbersome for packed curricula or too time-consuming for overworked faculty. One way to overcome this is to create viable authentic research experiences that can be integrated directly into the lecture-period and/or allow students to conduct research at home. As higher education begins to place a greater priority on quality online education, virtual CUREs can serve as a means to increase undergraduate participation in authentic research experiences and have already seen some success (22).

Here we report the implementation of a virtual CURE (vCURE) geared toward first-year undergraduates at Rutgers University Camden, a primarily undergraduate institution with a diverse and nontraditional STEM student population (90% commuters, 55% first-generation/low income, 28% African American, and 16% LatinX). The course allowed students to explore the intersection between the opioid epidemic and pain using the model organism *Drosophila* melanogaster at a cost of less than $15/student. Through self-report data, we find that this course improved science identity and key research competencies as assessed by the Undergraduate Research Student Self-Assessment (URSSA) survey (23, 24). This manuscript describes the structure and function of this 3-credit vCURE model and provides detail to adopt it in its entirety or adapt it to fit specific needs. Uniquely, our vCURE could be implemented in a hybrid manner as a project-based research experience that serves as the active-learning component of flipped classrooms ranging from Biology 101 to upper-level Neuroscience. We also describe details of tapping into the proliferation of virtual scientific conferences/webinars as a method to provide students with a unique training opportunity often missed by the financial constraints of attending in-person conferences. It’s our hope that this vCURE model could be widely and easily adopted for use nationally to increase access to authentic research experiences at any stage of the undergraduate curricula.

### Intended audience

The intended audience is first-year biology majors but is designed to run at any undergraduate level. Additionally, we also had a handful of non-biology majors, suggesting it could be effectively run for a range of undergraduates.

### Learning time

This is a semester-long project-based 3-credit biology elective across 15 weeks. However, individual components of the course can be adopted to fit into smaller time periods. Students can expect weekly to spend 1 hour watching pre-lecture videos, 3 hours in synchronous sessions, and 1-6 hours in asynchronous sessions working on the project or assignments.

### Prerequisite student knowledge

There is no prerequisite knowledge and students do not need experience working with *Drosophila* or in a lab setting.

### Learning objectives

1. Understand molecular mechanisms of addiction. (Comprehension)
2. Describe molecular mechanisms by which pain is governed. (Comprehension)
3. Demonstrate how to perform a search for primary literature. (Application)
4. Use *Drosophila melanogaster* laboratory techniques to address research questions. (Application)
5. Apply laboratory techniques to test hypotheses. (Application)
6. Generate a hypothesis based on primary literature. (Synthesis)
7. Critique scientific talks from a national conference. (Synthesis)
8. Support hypothesis with data. (Evaluation)

### Learning Outcomes

1. Describe neurological concepts of pain. (Comprehension)
2. Explain neurological concepts of addiction. (Comprehension)
3. Identify potential interest in scientific research. (Comprehension)
4. Compile scientific data to present to scientific audience. (Synthesis)
5. Formulate original hypothesis based on scientific literature. (Evaluation)

## Procedure

This course integrates experiential learning into a flipped-classroom environment by implementing *Drosophila*-based research into lecture periods. It is designed as a 3-credit 15-week course that meets twice/week but can be modified as the instructor sees fit to run in a shorter curriculum. While this course was run in and is ideal for a virtual setting, the original concept was developed for an in-person environment to place a research-based project into a traditional 3-credit flipped classroom without the need for including an additional co- requisite lab.

Each week is broken into two days: 1) content days where in-class discussions/activities focus on understanding required content and 2) lab meetings where in-class discussions/activities focus on completing a semester-long research project. Student assignments/activities are broken into “individual work” and “group work”. All work and activities are designed to mirror that which would be experienced in a traditional research lab.

Below are information/materials necessary to run this course within the scope of students learning about the neuroscience of addiction and chronic pain. However, the components can be used for other topics, and other protocols could be adopted to the content of interest. Thus, we have written this manuscript to facilitate faculty implementing the curriculum in its entirety or identifying individual components to adopt/modify/supplement their courses.

### Materials

Students will need the following:

1. Computer/internet access.
2. Video conferencing platform such as Zoom for synchronous meetings.
3. Cloud-based word processor/database/presentation programs such as those provided by Google Docs for collaborative work.
4. Communication platform for instant conversation/troubleshooting such as Slack.
5. Learning management software for classroom organization such as Canvas.
6. “Lab-in-a-box” that includes *Drosophila* and all necessary tools for students to carry out the behavioral experiments in the semester-long research project. This “lab-in-a-box” is described within the Lab Manual (Appendix 4) and Assembly Plan (Appendix 5).

### Student Instructions

Students are responsible for both individual and group work in the form of leading synchronous/asynchronous discussions, quizzes, presentations, and hands-on research activities. Detailed student instructions for each component can be found in the Course Syllabus (Appendix 1), Course schedule (Appendix 2), Journal club worksheet (Appendix 3), Lab Manual (Appendix 4), and Peer assessments (Appendix 6).

### Faculty Instructions

Below are instructions, commentary, and advice for the successful implementation of each component of the course. Further commentary on each component can be found in the Syllabus Annotations (Appendix 1).

#### Designing the Research Project

Faculty should design a semester-long research project that is easily approachable by students in the format of a traditional CURE. We recommend using backwards design that starts with a straight-forward research question/hypothesis/prediction and includes simple experimental design/variables. These can be constructed, as done here, by using *Drosophila* and simple behavioral assays. Once these are established, all other course content can be developed. We advise balancing impact with simplicity when designing this. For our project, we utilized four simple behavioral assays to study the impact of chronic pain on the development of addiction in *Drosophila*.

Given the limited in-class time, project plans should be fully developed prior to starting the course. See the Lab Manual for details on our project (Appendix 4). Faculty may feel free to use our project design to explore the same question, a similar question, build a different project based off the included behavioral assays, or develop an entirely different research project. We encourage student feedback when designing these variables but within reason to ensure the project has scientific authenticity and is not simply repeating existing published findings. For example, while we explored the impact of chronic pain on addiction, students may be interested in exploring how sleep, diet, or other factors impact addiction. Exploring an unknown instead of confirmatory experimentation instills greater buy-in from students and is the driving essence of authentic CUREs. We also suggest using the simple-to-approach behavioral assays. We used the negative geotaxis, sensitivity, tolerance, and Capillary Feeder (CAFE) assays to assess addiction. If built around simple and consistent behavioral assays as the dependent variable, faculty can explore a range of relevant topics as independent variables.

#### Conducting the Research Project

We include details for building a “lab-in-a-box” at the cost of approximately $15/student. If deviating from our project design, we recommend taking time to consider all necessary equipment, including gloves, paper towels, additional fly food, and vials. We also recommend waiting to send this “lab-in-a-box” until the experiments actually begin (approximately week 5) since flies need to be flipped to new vials regularly and students do not begin their experiments until week 5/6.

While students are broken into groups of 4-5, each student is responsible for collecting a single set of experimental data points for each assay (i.e., every student will conduct all experiments instead of a single group being assigned a single assay). Thus, a student should complete a set number of replicates, but each student’s data will be considered a single “n” to ensure variability of environment is accounted for, especially in virtual settings. This helps validate student results if the data are to be published. It is to be expected that not all students will produce sufficient data as all experimentation features potential methodological failures or impassable troubleshooting. Students should be reassured that this is a natural part of research and should not feel pressure to rush any experiments to preserve reliability of the data. Additionally, some students may wish to abstain from certain experimental procedures due to personal ethics or discomfort with flies. These students can virtually conduct the experiments with peers to still gain the underlying concepts and knowledge of the experiment.

As seen below, each group is assigned a different research presentation. To foster ownership and project management skillsets, these groups are responsible for coordinating data consolidation and analysis for their particular presentation. This may include groups creating a central repository with a cloud-based software for data upload.

#### Content Days

The first session of the week is a lecture period that includes active learning and traditional lectures that reviews essential content. Critically, the content should be identified with backward design framed around the goals of the overarching research project. For example, we identified critical knowledge necessary to understand the project (basic neuroscience, neuroanatomy of pain/addiction, *Drosophila* biology, etc.) and used backwards design to build the course content structure. These days include pre-lecture videos (Appendix 2), weekly quizzes, presentation of recent science news, and traditional lecture content.

#### Lab Meeting Days

The second session of the week includes group-based journal clubs, presentations, and technique instruction which can be seen in the Course Schedule (Appendix 2). The first few weeks reinforce concepts in experimental design, responsible conduct of research, basic statistical analysis, and presentation of data. The following weeks focus on group-led journal clubs related to the research project. Once students have received their “lab-in-a-box”, faculty use this time to instruct students on experimental techniques. We recorded these sessions for future reference and used break-out rooms for troubleshooting. Once students begin to collect data, these sessions are then used for the group-led research presentations.

### Suggestions for Determining Student Learning

Rubrics are included with all assignments. In addition to each rubric, two peer surveys are provided: 1) Peer Evaluation – group members rate each other’s involvement. 2) Presentation Peer Feedback – non-group members rate presenting group. Weekly quizzes are designed not only to test comprehension of the pre-lecture videos, but also provide incentive to prepare for class discussions.

### Sample data

We have provided examples of each of the four research presentations (proposal, RIP I, RIP II, and Thesis Defense), a Conference Debrief Presentation, a Journal Club Presentation, and a Journal Worksheet Example (Appendices 8-11).

### Safety issues

The procedures and contents were designed to comply with the American Society of Microbiology Guidelines for Biosafety in Teaching Laboratories. Although none of the items in this “lab-in-a-box” are hazardous, we recommend students attend an in-person or virtual lab safety training. We utilized an online CITI Right-To-Know lab safety training program. Of note, however, these boxes contain live *Drosophila*, low concentration ethanol (max 50% EtOH in comparison to hand sanitizer at 60% EtOH), and small glass capillary tubes.

## Discussion

### Field testing

This course was initially conceived as a strategy to integrate a low-cost CURE into a flipped classroom as an in-person semester-long project-based activity. Whereas flipped classrooms often contain a series of active learning exercises and CUREs are often their own separate entity, our curriculum can be plugged into a traditional flipped lecture-based course. This allows for the expansion of CUREs while avoiding the common impediments to introducing them more widely since any flipped course could rely on our CURE model for its active-learning component. As such, we built this CURE around a popular course that ran in two previous semesters at the 400-level called “The Neuroscience of the Opioid Epidemic”.

We initially piloted the virtual aspect of this CURE in the summer of 2020 when we partnered with the non-profit research hub “eCLOSE” that had developed a fully virtual bioscience research curriculum with a Drosophila-based “lab-in-a-box” for students ranging from middle school to college. During that summer, four Rutgers Camden undergraduates participated in this program along with 49 students from other institutions to pilot the “lab-in-a-box” curriculum and identify methods to adapt it for use with our pain/addiction course plan. We combined this virtual component with other mechanisms ideal for the lecture period that were developed and tested in our more traditional 300-level CURE; we then launched our lecture-period-based virtual CURE in the Fall of 2020. Thus, components of this virtual CURE have been field-tested over several semesters, piloted in the summer of 2020, and fully tested in a 100-level undergraduate course with 19 honors students during the Fall of 2020 (See demographics for the 12 who participated in the survey in Table 1).

**Table 1:** Student demographics. SES refers to Socioeconomic Status. Science Experience is the percentage of students involved in those associated activities related to science.

| Group |  | Demographics |  |  |  |  | Major |  |  | Year |  |  | Science Experience |  |  |  |  |
| --- | --- | --- | --- | --- | --- | --- | --- | --- | --- | --- | --- | --- | --- | --- | --- | --- | --- |
|  |  | n | Mean Age | Male | Female | Low SES | URM | Biology | Health Sciences | Other | 1st | 2nd | 3rd | Have previous research experience | Attended science-related museum or watched science-related documentary outside of classroom. | Had a conversation about a science-related topic with family or friends. | Read a science-related book or magazine outside of the classroom. |
| Full Cohort |  | 12 | 19.75 | 42% | 58% | 25% | 8% | 25% | 58% | 17% | 75% | 8% | 17% | 0% | 83% | 100% | 58% |
| Not-First-Gen |  | 6 | 18 | 67% | 33% | 0% | 0% | 17% | 67% | 17% | 83% | 17% | 0% | 0% | 83% | 100% | 50% |
| First-Gen |  | 6 | 21.5 | 17% | 83% | 50% | 17% | 33% | 50% | 17% | 67% | 0% | 33% | 0% | 83% | 100% | 67% |

### Evidence of student learning

Learning objectives 1-2 and outcomes 1-2 were evaluated with the formative assessments of weekly quizzes. Learning objectives 3 and 7 were assessed with journal club presentations, journal club worksheets and a final conference debrief presentation (Appendices 8, 10, 11). Notably, it was impressive to see the students digest and discuss research articles in this field as primarily first-year students. We strongly believe with guidance, non-honors students could also achieve similar outcomes. Learning objectives 5-6, 8 and outcomes 3-5 were assessed with the formative and summative assessments of research presentations (Appendix 9). For these presentations, students must be able to conduct the experimental assays and collaborate with their peers outside their group to consolidate, analyze, and present their data.

We additionally used two validated survey instruments in a pre-test/post-test design to assess the effectiveness of our vCURE on several science-related outcomes. The pre-test took place at the end of the fourth week of class before students received their “lab-in-a-box”, but after they had a month to gain familiarity with the study and content. This allowed us to examine the effects of the course’s research component. 12 of the 19 students took part in both the pre- and post-tests. We collected demographic data indicating first-generation college student status, major, age, gender, and year (Table 1). To ensure these students were well matched, we also confirmed that they did not have any previous research experience. However, we note their interest in science given their participation in a range of extracurricular science-related activities (Table 1).

We first assessed science identity which is a measure of how much a student identifies as a scientist or science trainee (23). This measure positively correlates with success, academic retention, and whether the student enters a science occupation. On a 1-7 Likert scale, science identity significantly increased from 4.75 ± 1.36 to 5.33 ± 0.89 in the full cohort (Figure 1A, p = 0.0463). We saw no differences when comparing first-generation to non-first-generation students. Interestingly, science identity was significantly less in females than males at both the pre- and post-timepoints (Figure 1B, C, p < 0.05 and < 0.001 respectively).

**Figure 1:**
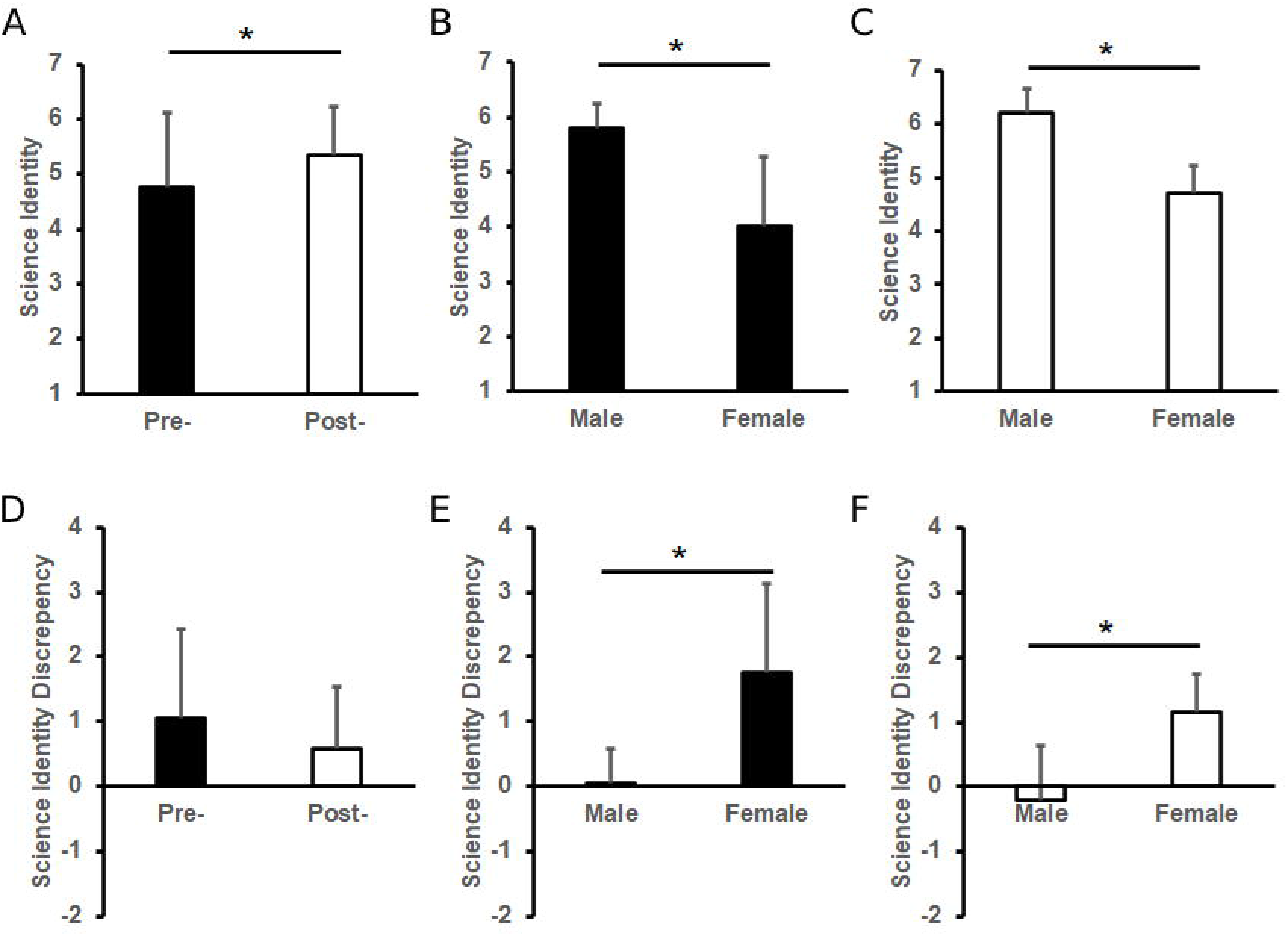
Science Identity Measures. Science Identity of full cohort at pre- and post-time points (A), male vs female at pre- timepoint (B), and male vs female at post- timepoint (C). Science Identity Discrepancy of full cohort (D), male vs female at pre- timepoint (E), and male vs female at post- timepoint (F). Pre- timepoint assessments occurred during week 4 of the course prior to receiving the “lab-in-a-box” while post- timepoint assessments occurred during week 15 at the end of the experimentation period. n = 12 for full cohort (7 females, 5 males) with * indicating p < 0.05.

We also measured science identity discrepancy which assesses the difference between how a student perceives themselves in relation to how they perceive others see them in science. This comparison creates a scale where more positive numbers indicate a student rates themselves less than they think others would rate them in science. Thus, positive values could be interpreted as a measure of “science imposter syndrome” while negative values could be interpreted as a measure of “science underdog status”. While no changes were seen in the full cohort comparing pre- to post-, we did see gender differences at each time point (Figure 1D). At both the pre- and post-timepoints, males had neutral science discrepancies (Figure 1E). Females, however, had greater discrepancy scores than males at both time points, suggesting greater levels of imposter syndrome (Figure 1F). There were no differences between first-generation and non-first-generation students.

We then used the URSSA survey to evaluate whether our vCURE was a successful program (24). This validated 34-question survey reports four critical research measures: Thinking and Working Like a Scientist, Personal Scientific Gains, Scientific Skills, and Researcher Attitudes/Behaviors. We found significant increases in all four measures across the total cohort (Figure 2) but no differences between first-gen and non-first-gen students or between genders. Notably, we find no differences amongst any of the groups in regard to their baseline research confidence which is a measure of self-efficacy to perform science-related tasks (Table 2). Combined with the gender differences in science identity discrepancy, we posit that while female students within our class may feel less capable than those around them, a quantified self-assessment of their own ability suggests otherwise.

**Table 2:**
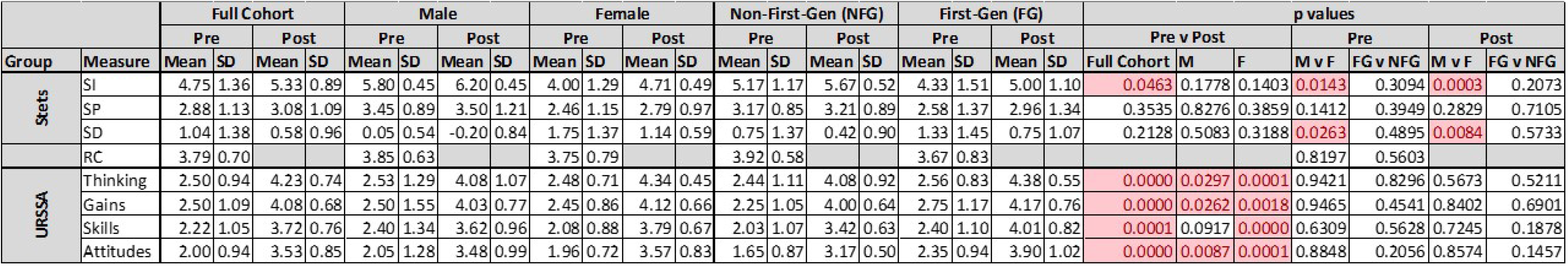
Means ± standard deviation for all survey instrument measurements and p values for all comparisons. STETS refers to the Science Identity questionnaire developed by Stets et al. in 2017 that measures Science Identity (SI), Science Identity Prominence (SP), and Science Identity Discrepancy (SD). RC indicates the Research Confidence Questionnaire. URSSA is the Undergraduate Research Student Self-Assessment Measures developed by Weston et al. in 2015 that measures thinking and working like a scientist, personal scientific gains, improvement of scientific skills, and improvement of researcher attitudes/behaviors.

**Figure 2:**
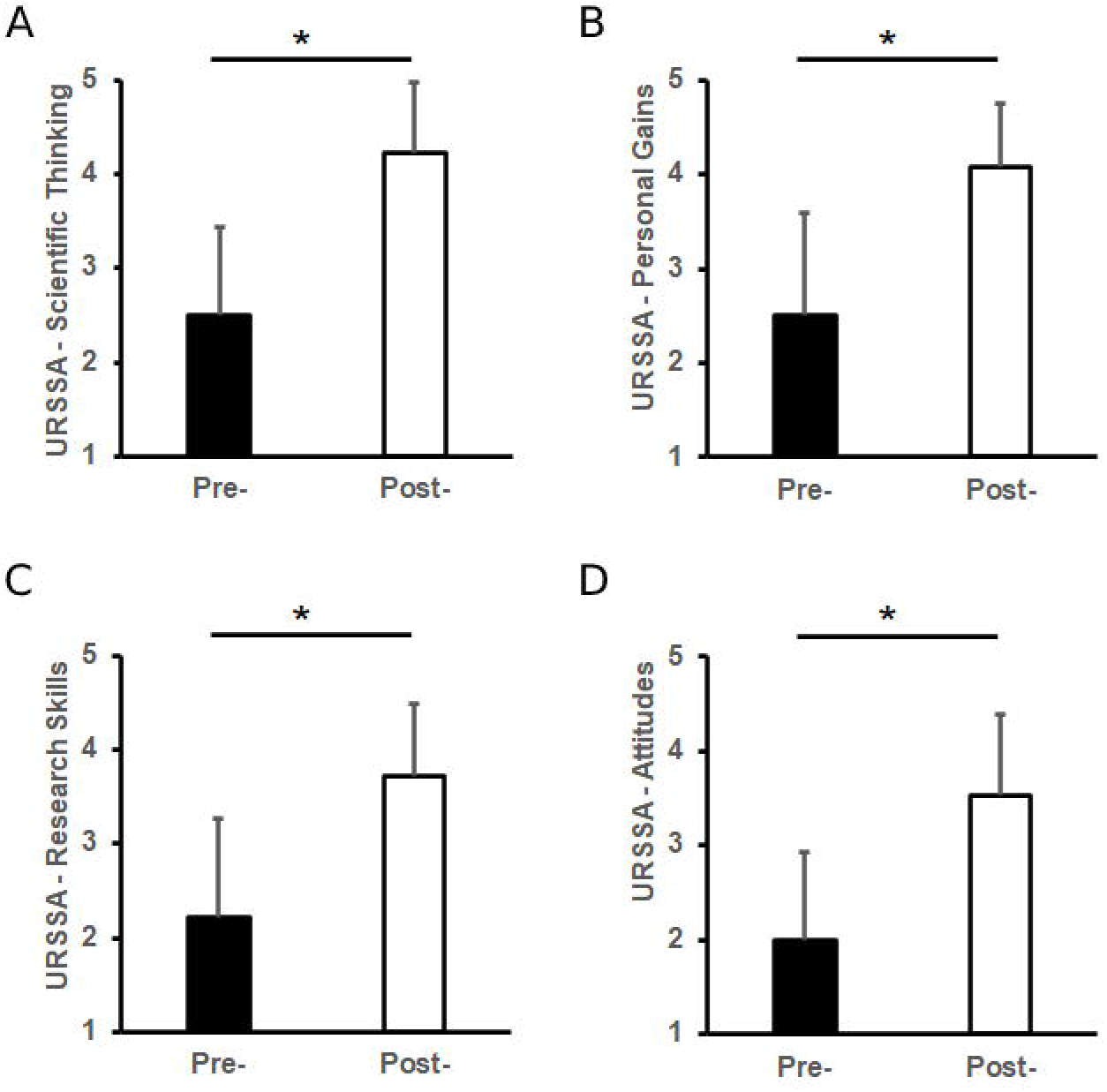
URSSA Scores. Scores from the Undergraduate Research Student Self-Assessment for the full cohort: Thinking and Working Like a Scientist (A), Personal Scientific Gains (B), Scientific Skills (C), and Researcher Attitudes/Behaviors (D). Pre- timepoint assessments occurred during week 4 of the course prior to receiving the “lab-in-a-box” while post- timepoint assessments occurred during week 15 at the end of the experimentation period. n = 12 for full cohort with * indicating p < 0.05.

These gender differences are not necessarily surprising since females in several fields indicate greater imposter syndrome than males (25, 26). However, it reinforces the importance of still pursuing systemic improvements that encourage women in STEM regardless that the National Institutes of Health no longer considers women as underrepresented in the biomedical sciences. While we didn’t see any differences in first-generation students, the URSSA assessments may instead suggest that this vCURE can be effective regardless of familial college experience.

### Possible modifications

With the inevitable proliferation of flipped classrooms after COVID-19 forced most biology faculty to pre-record their lectures for the first time, we believe there is an unprecedented opportunity for integrating CUREs directly into the lecture-period as the active learning component of a flipped classroom. Given that our CURE could be run in either a virtual or in-person capacity, we are confident this model could be widely adopted. Further, the recent comfort with virtual settings could expand the accessibility of research experiences by adopting this vCURE into a range of settings such summer bridges or REUs (Research Experiences for Undergraduates) without the need for expansive infrastructure or large budgets.

While we ran this as a full 3-credit 15-week course, it could be consolidated to a shorter period by removing experiments. Combined with its low-cost and virtual capacity, it could be integrated into non-curricular activities such as summer bridge programs. This vCURE could be a low-cost method to foster science identity prior to a student entering their first college-level biology course. It could also easily be scaled up to large entry-level biology courses which have traditionally found it challenging to integrate active learning. Further, these activities are beneficial for students at any level and thus could be used in upper-level courses.

Our model can be modified to function in either a virtual or in-person setting and could be connected to any scaffolded biology course. The course could also be adopted in its entirety or specific components integrated into other curricula. Further, the behavioral assays could be adopted for research studies exploring a wide range of research topics outside the realm of pain and addiction.

One of the defining characteristics of immersing oneself fully into the scientific community is interacting with peers and colleagues at scientific conferences. This essential experience is often restrictive to undergraduate students either due to financial reasons constraints of travel or registering. As conferences begin to move back to in-person settings, we are confident webinars and virtual seminars will continue to be accessible for students to tap into within the structure of our vCURE.

## Acknowledgements

We thank the many Rutgers Camden undergraduates who provided their feedback as we developed and launched this course. We thank Dr. Kwangwon Lee who provided mentorship and guidance to make this course and manuscript a success. We also thank the Rutgers Camden Honors Program and Provost Office for funding the “lab-in-a-box” and the NIH K12 IRACDA PennPORT program (K12-GM081259) for funding the virtual conference travel. Finally, we thank the American Society for Cell Biology (ASCB) Promoting Active Learning and Mentoring (PALM) Network for their funding that allowed the authors to develop this course.

## Supplemental Materials

### Student Documents

<u>Appendix 1</u>: Course Syllabus - This course syllabus includes annotations for faculty within the comments to understand how to implement each portion of the course.

<u>Appendix 2</u>: Course Schedule - This document provides a list of all topics and recommended timeline for a 15-week course. Annotations for faculty within the comments are included for further guidance.

<u>Appendix 3</u>: Journal Club Worksheet - This worksheet can be converted into an LMS quiz for faster grading and easier submission.

<u>Appendix 4</u>: Lab Manual - This lab manual includes the framing of the semester-long project, details on the “lab-in-a-box”, and all protocols necessary to carry out the experiments.

### Faculty Documents

<u>Appendix 5</u>: “Lab-in-a-box” Assembly Plan - This document includes the assembly instructions, items, and details for where and how to purchase the items.

<u>Appendix 6</u>: Peer Evaluation questions - This document includes questions for peer evaluations given twice in the semester. It is advised this document be uploaded to an online quiz system such as Google Forms.

<u>Appendix 7</u>: Research Presentation Rubrics - This document includes the rubrics for all research presentations.

### Example Student Work

<u>Appendix 8</u>: Journal Club Worksheet Example – this is an example of a student’s journal club worksheet.

<u>Appendix 9</u>: Research Presentations - This document includes the Research Proposal, Research in Progress I/II, and Thesis Defense Presentations.

<u>Appendix 10</u>: Journal Club Presentation – This is an example of a journal club presentation given by one group.

<u>Appendix 11</u>: Conference Debrief Presentations - This document includes an example presentation for the conference debrief assignment.

### Other

<u>Appendix 12</u>: Survey Instrument Design and Details

## Notes

**Sources of Support**: NIH K12 IRACDA PennPORT program (K12-GM081259) provided funds for conference registration. American Society for Cell Biology (ASCB) Promoting Active Learning and Mentoring (PALM) Network provided funds for collaboration between authors. Rutgers Camden Honors Program and Provost Office provided funds for the “lab-in-a-box”.

**Conflict of Interest Notification**: The corresponding author, Nathan T. Fried, and Edward A. Waddell declare no potential conflicts of interest with respect to the research, authorship, and/or publication of this article. Dara Ruiz-Whalen and Alana M. O’Reilly declare that they are the Chief Learning and Chief Scientific Officers, respectively, for the Philadelphia-based non-profit organization, eCLOSE Institute.

### Competing Interest Statement

The corresponding author, Nathan T. Fried, and Edward A. Waddell declare no potential conflicts of interest with respect to the research, authorship, and/or publication of this article. Dara Ruiz-Whalen and Alana M. O'Reilly declare that they are the Chief Learning and Chief Scientific Officers, respectively, for the Philadelphia-based non-profit organization, eCLOSE Institute.

