## Supplemental Files for "Flying in the Face of Adversity: A Drosophila-based Virtual CURE Provides Semester-long Authentic Research Opportunity to the Flipped Classroom"

### Appendix 1 – Annotated Syllabus

#### NEUROSCIENCE OF THE OPIOID EPIDEMIC

#### COURSE DETAILS

**Course Description:**

Time: Monday/Wednesday 2:05-3:25
Virtual Location: Neuro Zoom Room (you can gain access in canvas)

This special honors college seminar course will approach the opioid epidemic from a basic science perspective. The course will cover the basic neuroanatomy, neurophysiology, and neuropharmacology of sensory systems and addiction. Students will first study how sensory systems work with a focus on pain sensation and then study drugs of abuse and addiction with a focus on opioids. The class will explore the interplay between chronic pain and the opioid epidemic to approach this public health issue from a basic science perspective.

Class will include a combination of discussion-based lectures, student presentations, a unique semester-long authentic research project using fruit flies conducted virtually (yes, we'll send you the necessary equipment and flies!), and participation in a national scientific conference that will be virtual this year. Ultimately, there is a mixture of individual and group work and you should all view each other as members of a large lab working together on a single research goal.

Uniquely, this course will be taught by two neuroscientists who have a strong focus on both research and education. Dr. Fried is at Rutgers-Camden and trained in a special postdoctoral research/education program at the University of Pennsylvania called IRACDA. Dr. Waddell is currently an IRACDA fellow. Both have a strong dedication to breaking down barriers in science for students of all levels and backgrounds, a drive they both developed coming from first-generation and/or low-income backgrounds. Together, they hope this class can give you a taste of what it's like to be a scientist.

**Professors:**

Nathan T. Fried, PhD (email address)
Assistant Teaching Professor, Department of Biology, Rutgers Camden
Research Area: Neuroscience, Pain, Opioids, Sleep, Biology Education
Office Hours: M (11am-1pm) & W (3:30-5:30pm) Virtual Office | Bookings Page (you can pop in during office hours, but an appointment blocks YOUR time)

Edward Waddell, PhD (email address)
NIH IRACDA PennPORT Postdoctoral Fellow, Department of Neuroscience, University of Pennsylvania
Research Area: Neuroscience, Genetics, Molecular Biology, Developmental Biology, Biology Education
Office Hours: M (3:30-5:00pm) & W (11am-1pm) Virtual Office

**Note from the professors:**

It is our hope to create a class that is challenging, informative, and enjoyable. The topics in this course, however, may affect some students more than others since the opioid epidemic is widely experienced by so many of us in different ways throughout the Philadelphia area. Given the stigma behind some of these topics (whether it be the personal experience of chronic pain or addiction), we ask all students to be respectful of one another's experiences to create a safe atmosphere to engage in these challenging topics.

**Commented [nf1]:** this can be any topic that includes drosophila behavioral techniques or easy to adopt tools.

**Commented [nf2]:** link to the virtual space you'll use.

**Commented [nf3]:** Link to an Outlook Bookings page reduces emails to set up appointments.

Virtual office hours increases the numbers of students who actually use office hours because its easier to tune in.

Further, given COVID turning this course into a virtual experience, it's paramount that you, as the student, are as engaged as possible. We will be understanding of missed classes, distractions at home, or other challenges, but the class will only work if you throw yourself into the material. We're also always open to your suggestions on how to better run the course, so be vocal about your feedback! We're here to listen and improve your experience. So, please, be engaged, speak up, speak out, and let's make this semester a memorable one.

**Learning Objectives:**

Understand the molecular mechanisms of addiction. (Comprehension)
Describe the molecular mechanisms by which pain is governed. (Comprehension)
Demonstrate how to perform a search for primary literature. (Application)
Use laboratory techniques specific for working with *Drosophila melanogaster* to address research questions. (Application)
Apply laboratory techniques to test hypotheses. (Application)
Generate a hypothesis based on primary literature. (Synthesis)
Critique selected scientific talks from a national conference. (Synthesis)
Support hypothesis with data. (Evaluation)

**Learning Outcomes:**

Describe the neurological concepts of pain. (Comprehension)
Explain the neurological concepts of addiction. (Comprehension)
Identify potential interest in scientific research. (Comprehension)
Compile scientific data to present to a scientific audience and peers. (Synthesis)
Formulate an original hypothesis based on scientific literature. (Evaluation)

**Note regarding learning objectives and outcomes:**

It's important to understand the difference between expected learning objectives and expected learning outcomes of this course. The learning objectives serve as a means to organize the accomplishment of certain tasks in order to progress throughout this course. The learning outcomes instead represent the end goals of this course (what we want and hope that you take with you after the course ends). The outcomes of this course are considered to be goal-directed, meaning that they are designed with **YOUR** goals in mind. By taking this course, it can be assumed that you have an interest in Neurobiology, whether that be clinical or laboratory. Therefore, the outcomes of this course highlight critical skills and knowledge that are essential for your future studies and training in this field.

#### COURSE MATERIALS

This course does not have a textbook. Instead, we will use online resources. All class sessions will be recorded and put online.

<u>Excellent online guide to pain pathways information</u>:
http://nba.uth.tmc.edu/neuroscience/s2/chapter07.html

<u>Pain as an Art Form</u>: https://well.blogs.nytimes.com/2008/04/22/pain-as-an-art-form/?_r=0

Neuroscience Virtual Textbook (Free): https://nba.uth.tmc.edu/neuroscience/

Zoom: We'll use Zoom to meet.

Canvas: We'll organize the class in canvas.

Computer: You should have a computer and reliable internet access. If you don't, please let me know.

#### CLASS STRUCTURE

See excel sheet for each week's topic/plan/due dates. Below is an example of how each week will run.

Weekend: Students watch pre-module videos, complete the week's 5 question quiz, and complete their journal club worksheet (if due for the module) before coming to class.

Mondays (content days): Student-led News story (2:05 - 2:15 pm)
Break-out room conversations (2:15 - 2:25 pm)
Lecture (2:25 – 3:25 pm)

Wednesdays (lab meeting days): Journal Club or Presentations or Research Demos

**Commented [nf4]:** The remaining time in the content day can be used to reinforce core concepts with traditional lecture or active learning as the faculty see fit. Critically, these lectures are meant to reinforce core concepts in a similar fashion one might explain scientific concepts to a new trainee in a lab as opposed to a comprehensive survey of all components of a topic. Using backwards design, faculty should focus on what students need to know to understand the research project as opposed to a survey of information in preparation for exams, which are not included in this course.

#### COURSE ASSIGNMENTS, ACTIVITIES, & DETAILS

This class is designed to be hands-on. As such, there are several activities students will participate in. All of these activities will help us finish our research study together as a collective lab.

**Individual Work**: Some work will be conducted on your own.

**Pre-Lecture Videos:** We have curated a list of videos that are interesting and enjoyable to watch. You'll be responsible for watching these videos before coming to class. They will let you dip your toes into the week's content.

**Commented [nf5]:** Depending on the week's content topic, faculty should curate a short list of online videos that expose students to the week's content without overburdening student time since some weeks they will be working on several other projects. We suggest short youtube videos that introduce students to the week's topics. Collectively, the videos should not be longer than 45 minutes. We have included all pre-lecture videos in Appendix 2.

**Weekly Quizzes (due before first class of the week):** Each week, there will be a 5-question quiz you can answer from the pre-lecture videos or other resources on the internet. Some questions will be harder than others and will require you to do some digging. You will have the entire previous week to take the quiz and it will not be timed.

**Commented [nf6]:** To incentivize watching the videos, a short untimed 5-question quiz was due prior to the beginning of class. This ensures students are ready to participate in discussion.

**In the News Curator of the Week:** It's valuable to stay current in your understanding of any field. This weekly exercise will facilitate this by having you each responsible for learning about recent discoveries related to the week's topics. Each week, 1-2 students will be assigned as a "curator of the week" where they will participate in/lead the following activities.

**Commented [nf7]:** To foster conversation, reinforce key concepts, and encourage independent learning, students will engage with recent science news related to the week's topics and research project.

Monday News Presentation: Each Monday, the "curator of the week" will present a 5-7 minute PowerPoint (or google slides) presentation about a news story from www.The-Scientist.com related to the week's topic.

**Commented [nf8]:** One student a week will identify a news story related to the week's content at www.The-Scientist.com and give a 5-minute presentation at the beginning of class. Each presentation should end with a student-generated question for reflection in the breakout room conversations. We use The-Scientist.com since it is a well-known daily publication on a range of science topics. We also want to do this to familiarize students with the website so they can become more used to reading science news from a reputable news outlet.

Grading Rubric:

- Show us the news story and explain what research was conducted. Try to identify the a) field, b) hypothesis, c) results, and d) how this advances the field. The news story must be from www.The-Scientist.com. (50%)
- Explain why you chose the news story for the week's topic. (20%)
- Include 1-2 questions at the end of the presentation to prompt conversation for the class. (20%)
- Presentation is between 5-7 minutes long (10%)

Canvas Discussion Moderator (due Monday morning): The "curator of the week" must make a post in Canvas Discussion that includes the link to the article they presented on along with the questions they used to generate conversation and an art piece they find on the internet that is somehow related to the week's topic, along with a) the artist's name and b) the reason you chose the art piece. You should monitor the class's comments on the posts and try to engage your peers.

**In the News Online Conversation:** What's the point in posting things if it doesn't generate conversation, right? Each week's Curator will post their news story to Canvas. You will each be required to contribute to the conversation on Canvas and should be active at least 70% of the semester's weeks. Although this will be monitored for your activity, the goal is to generate insightful and meaningful conversation, not just busy work.

**Journal Club Worksheet (due night before journal club days):** If you are not presenting a journal club for the week, you should fill out a journal club worksheet. For this class to work, it is ***IMPERATIVE*** that you have read the journal club assigned for the week. Part of this worksheet includes preparing two questions/points to ask the presenter. At the time of the presentation, students are encouraged to ask these questions and participate in the discussion. Consider this a safe place for practicing having the courage to ask questions!

**Drosophila Research Study:** You will each receive a vial of drosophila to conduct a series of experiments that will test the effects of chronic pain on behavior and addiction. You are expected to participate in the demos and run your experiments independently on your own time. You will not be graded on this, but the class presentations rely on you completing the research and providing your results.

**Group Work**: We will pre-assign 4 groups of 5 students to complete various assignments. Each member of the group is required to equally contribute to both the oral presentation and presentation preparation. You should create a group name.

**Monday Break-out Rooms**: The break-out rooms will not be graded, but they're an opportunity to "prime yourselves" to be active learners, instead of passive ones, regarding the week's material. During this brief time, you'll be able to chat with your group about the "arts and sciences" presentation or the day's lecture material. Discuss anything you don't understand so that you're ready to discuss them with us during the lecture.

**Journal Club Presentation:** Student groups will prepare a 30-minute PowerPoint presentation on an assigned research article. Each group will present once. These papers will highlight an active area of research on topics discussed in class. In preparation, students are encouraged to discuss the papers, as well as any issues related to their presentation, with the instructors in advance. For reference, when we give journal clubs in our own lab meetings, it takes us about 5 hours to read, distill down the info, and make nice slides. At your early stage in your research career, it can take much longer. So, don't delay getting started.

A lot of this is going to be above your head because many of you are first-year students who have never read research articles. But give it your best shot. We'll grade fairly. Your job as the presenter for the week

**Commented [nf9]:** To encourage classroom-wide conversation related to the week's news story, the presenting student will post the news story on the LMS and will moderate peer conversation

**Commented [nf10]:** To encourage participation in the conversation, all students are required to respond to a minimum number of posts throughout the semester.

**Commented [nf11]:** To ensure all students have read the paper prior to the presentation, any non-presenting student will fill out a worksheet that distills down the important aspects of the paper.

The goal is not for them to be experts on this paper, but instead to be familiar enough with the paper to understand the presenting groups.

This worksheet includes manuscript meta information to provide students, especially first-generation students, with experience understanding how journals work and what universities conduct this type of research. The Worksheet is in Appendix 3 and an example from a student is in Appendix 8.

**Commented [nf12]:** To provide an opportunity for group engagement, students will briefly meet with their groups to discuss the week's content and news story question of reflection. This time is important for developing solidarity and camaraderie within the group.

**Commented [nf13]:** Journal clubs are traditionally given in research labs where one person presents the details of a research article that the rest of the lab has read. This keeps a lab up to date on current literature in their field.

To increase familiarity with the research project field, each group will present one research article related to the project in the format of a traditional journal club. Each group will present once in the semester and thus faculty should identify an equal number of critical papers to the number of groups within the course. Students are encouraged to present in such a way that helps their peers understand the material.

Example presentations are in Appendix 10.

is to essentially teach your peers about the research article. You are the expert on the paper and thus, your goal is to guide us through it so that we all understand it.

**Structure of Journal Club Presentation/Rubric (remember, ~2 min for every slide):**

Introduction/Background (~10 min) (50%): It is VITAL to frame your presentation of the article. Use their own introduction as your guide. Put their findings into context so that once you delve into the data, the audience will have all the information they need to actually understand the data.

Describe the following in your intro:
1) The field the authors are exploring.
2) What is not known about the field.
3) The question they are trying to answer with their research, review, or theoretical paper.
4) How they propose to answer or approach the question.

Where does it fit? (~2 min) (10%): Now, discuss how this paper fits into the context of the course.

Results (~10 min) (10%): You should go figure by figure and describe the logic for doing that experiment, the way the experiment works (i.e., the methods used), and what the conclusion is from each figure. This will let you move figure to figure smoothly. Consider the data as a story. Tell us the story.

Conclusions (~2 min) (20%): Now, take that data or theoretical framework and put it into context with what you presented during the intro. Tell us how it solved the question they proposed in the intro.

Questions/Discussion (~6 min) (5%): Be ready to answer any questions from the audience. It is OK to say, “I do not know.” Being a scientist means being comfortable with the unknown. You should also prepare your own intellectual questions to ask the audience to generate conversation.

Class Score (5%): Receiving critical feedback from your peers is essential to become a stronger presenter. All students in the class will give you critiques and a final score for your presentation. Please see provided rubric for details on the assessment.

**Journal Club Moderators**: One group will be assigned to be the moderator for the week’s journal club. Each group will be the moderator once. It’s their job to keep the conversation going and active. Your group should prepare at least 3 questions to ask the presenters.

**Commented [nf14]:** To increase active discussion during the journal club, each group will be assigned once to be the moderators for the day. They will field questions from the audience and be prepared to ask their own questions of the presenters.

**Research Project Presentations:** When a scientist conducts a years-long study, they have to present several times. The first presentation focuses on presenting the research idea to obtain feedback on it before it's started. Then there are several project updates where data is presented. And finally, once the conclusions are drawn, it's presented to the broader community where the audience tries to pick apart the conclusions and the scientist needs to defend their work. Each group will present one of these presentations. It'll require everyone working together, sharing their data, etc. Each group will be assigned to present **one of the following four presentations**. Each presentation is designed to build off each other, therefore by the end of the course, we’ll have a full series of presentations that was created by all course students. Students not presenting are expected to ask questions during the presentations. This not only builds your skills in asking relevant, thought provoking questions, but also allows each presenter to build their skills in thinking critically about what they are presenting.

**Commented [nf15]:** Traditional research projects generally include multiple iterative presentations throughout the completion of the work. To mirror this in the classroom, each group will be required to give one of these iterative talks. Each successive group should build off the presentation of the previous group throughout the semester as the research project is carried out. Students should view all groups working together on the common goal of completing this project, but each presentation will have different goals associated with it.

Example Research Project Presentations are in Appendix 9.

Grading Rubric**:** Each presentation will be different, but you can see the "goal" of each below. We leave it to your group's discretion and creativity to decide how you'll achieve that. Your grade will be dictated 90% by faculty assessment of whether you achieved that goal and 10% by whether the class believes you achieved the goal. Please see provided rubric for details on the assessment.

**Commented [nf16]:** Grading rubrics are in Appendix 7.

**Commented [nf17]:** The rationale for this evaluation is to use peer feedback as a means for students to help improve non-group members' presentation, critical thinking, and question answering skills. Additionally, through providing feedback to non-group members, each student has the opportunity to reflect on their own presentation and critical thinking skills.

**Proposal Presentation:** In this presentation, students are expected to present a thorough background of the course's research project including the scientific rationale for the research question, hypothesis (or hypotheses), and all relevant background knowledge needed to understand the project. Additionally, students are expected to outline the research project including a timeline of experiments and what research question each experiment is designed to address.

**Commented [nf18]:** One group will present the semester-long project as a research proposal, including the research question, goal, hypothesis, and aims.

Goal: Fully describe why and how the project is being pursued in such a way that stimulates interest in the work.

**Research in Progress I:** Students will present the class-aggregated results from the climbing and sensitivity assays. This will require collecting data from each class member's experiments, aggregating it, analyzing it, and making a conclusion based off it. Students will describe the experimental design, the scientific rationale for each specific experiment, and the results (in an appropriate graph or chart). Additionally, students should provide a brief reintroduction to the work including their scientific background, question and hypothesis (or hypotheses).

**Commented [nf19]:** Groups assigned to these presentations will present the results collected from the class in the format of a research in progress. These groups should coordinate with the rest of the class to aggregate, analyze, and distill the results and conclusions. Depending on the number of groups, these RIPs should be scheduled according to the research being conducted. We have two RIPs because we have four groups. This process of coordination builds independence and problem solving when trying to do project management.

Goal: Provide an update on the first set of results and determine a conclusion for us to make decisions on the next steps of the project.

**Research in Progress II:** Students will present the class-aggregated results from the tolerance and preference assays. This will require collecting data from each class member's experiments, aggregating it, analyzing it, and making a conclusion based off it. Students will describe the experimental design, the scientific rationale for each specific experiment, and the results (in an appropriate graph or chart). Additionally, students should provide a brief reintroduction to the work including their scientific background, question and hypothesis (or hypotheses).

Goal: Provide an update on the second set of results and determine a conclusion for us to make final conclusions on the project.

**Thesis Defense Presentation**: Students will create a final presentation about what these experiments conclude. Students are expected to present all data collected throughout the term. Students will describe the experimental design, the scientific rationale for each specific experiment, and the results (in an appropriate graph or chart). Additionally, students should provide a brief reintroduction to the work including their scientific question and hypothesis (or hypotheses). Finally, students will highlight the major conclusions of their work and potential future directions and implications of their work.

**Commented [nf20]:** One group will present the final project presentation in the format of a thesis defense.

Goal: Describe the full picture of the project, it's conclusions, and future directions so we can assess where to go after this class is over.

**Conference Debrief Presentation (During Finals Week):** A unique opportunity within this course is that each student will receive a one-year undergraduate membership to the American Society of Cell Biology (ASCB). As a part of this, each student will be expected to attend the virtual ASCB conference this year from December 2-16. Each group of students are responsible for selecting one talk to present to the class. To accomplish this, students will need to view the conference schedule and select a single talk they would

**Commented [nf21]:** If the faculty member has included virtual conference attendance for this course, each group will be assigned a talk or poster related to the project and will be responsible for presenting it to the rest of the class at the end of the semester. This will reflect activities seen in traditional research labs where the attending lab-mate discusses any recent findings relevant to the lab. We ran this during our final exam period.

Example Conference Debrief Presentation is in Appendix 11.

like to present on. We encourage you all to attend several talks, however, so you can learn and explore! We'll curate a recommended schedule for you once the agenda is released.

**Structure of Conference Debrief Presentation/Rubric (remember, ~2 min for every slide):**

Introduction/Background (~10 min) (50%): It is VITAL to frame the narrative of the presentation. Use their own introduction as your guide but add information as needed. Put their findings into context so that once you delve into the data, the audience will have all the information they need to actually understand the data.

Describe the following in your intro:
1) The field the presenter is exploring.
2) What is not known about the field.
3) The question(s) they are trying to answer with their research.
4) How they propose to answer or approach the question(s).

Where does it fit? (~2 min) (10%): Now, discuss how this presentation fits into the context of the course material.

Results (~10 min) (10%): You should go major experiment by experiment and describe the logic for doing that experiment, the way the experiment works (i.e., the methods used), and what the conclusion is from each. For figures, search the authors name using a literature search to see if their work is published. If there are no figures available, you should present their logical flow. Consider the data as a story. Tell us the story.

Conclusions (~2 min) (20%): Now, take that data or theoretical framework and put it into context with what you presented during the intro. Tell us how it solved the question(s) they proposed.

Questions/Discussion (~6 min) (10%): Be ready to answer any questions from the audience. It is OK to say, "I do not know." Being a scientist means being comfortable with the unknown. Also, you should prepare your own intellectual questions to ask the audience to generate conversation.

**Peer Evaluation**: You will have the opportunity to evaluate your group members twice (middle of course and end of course). The first evaluation will help you improve your team-work and the second will be a final evaluation of your participation in the group. Your grade will be an average of all your group member's assessment of you.

**Commented [nf22]:** The rationale for this evaluation is to provide students an opportunity to offer peer feedback to their own group members in an anonymous format. Having this evaluation as a part of a student's overall grade also encourages each student to contribute meaningfully to their group's work.

#### Calculation of Final Grades

Individual Work

| | |
|---|---|
| Weekly Quizzes (1 pts each x 14) | 14 pts |
| Monday News Presentation | 10 pts |
| Canvas Forum Moderator | 10 pts |
| Arts & Sciences Online Conversation | 10 pts |
| Journal Club Worksheet (2 pts each x 3) | 6 pts |

Group Work

| | |
|---|---|
| Journal Club Presentation | 10 pts |
| Journal Club Moderator | 10 pts |
| Research Project Presentation | 10 pts |
| Conference Debrief Presentation | 10 pts |
| First Peer Evaluation | 3 pts |
| Second Peer Evaluation | 7 pts |
| Total | 100 pts |

#### Classroom Policies

**Microphone and Video Policy:** When we can put a name to a face, it can be really helpful for all of us to connect virtually. We therefore strongly encourage you to have your video on. For your privacy, you can use a virtual background in Zoom. However, it is ultimately your choice whether to have your video on or not and we will respect that choice. Additionally, please keep your microphone off when not speaking. Background noise can obscure the conversation.

**Mental Health Statement:** Simply put, college and life are stressful and hard. The demands on you are immense, especially for those balancing a part-time job. If you are struggling or need someone to talk with, please reach out to me or visit the Rutgers Camden Student Wellness Center. We are here to help.

**Attendance/Tardiness**: We will not take attendance, but the course only works well if everyone participates. During these unprecedented times, we will have to work through days of unreliable internet access and family or loved ones who may be in challenging situations or sick/affected with COVID19. There is no need to alert us to any absences unless you are scheduled to present.

**Students with Disabilities Statement:** Rutgers University welcomes students with disabilities into all of the University's educational programs. To receive consideration for reasonable accommodations, a student with a disability must contact the appropriate disability services office at the campus where you are officially enrolled, participate in an intake interview, and provide documentation: https://ods.rutgers.edu/students/documentation-guidelines. If the documentation supports your request for reasonable accommodations, your campus's disability services office will provide you with a Letter of Accommodations. Please share this letter with us to discuss the accommodations needed as early in your course as possible. To begin this process, please complete the Registration form here.

**University Academic Integrity Statement:** Rutgers University takes academic dishonesty very seriously. By enrolling in this course, you assume responsibility for familiarizing yourself with the Academic Integrity Policy and the possible penalties (including suspension and expulsion) for violating the policy. As per the policy, all suspected violations will be reported to the Office of Community Standards. If in doubt, please consult the instructor and review the Academic Integrity Policy.

**Office hours policy:** Please **don't email me** to tell me you're coming to office hours. I get so many emails so it'll be hard to keep track of it. Instead, just use the bookings link I have at the beginning of this syllabus to book a 20-minute chunk (or more!) of time to meet with me. Booking helps me know if someone is coming so I don't walk away from my computer and to ensure other students aren't eating into your time. These office hours are time for me to chat with you about anything under the sun. Use them liberally. They're your time!!!

**Dr. Fried's Email policy**: I get a lot of emails every day. I teach all day/night on Mon and Wed so I won't respond to your email until Tues or Thurs (usually within 24-48 hrs). But if you haven't received a response, feel free to "ping me" again. Sometimes the flood of emails might bury yours. Pinging the message again is common in academia. Always feel free to remind someone of your email. Often, if they don't respond, it's just because they missed it; not because they are ignoring you.

**Dr. Waddell's Office hours policy:** If you know that you want to meet with me during my office hour periods, please email me ahead of time so that I can prepare for our discussion. My office hours will be open access; however, I will give priority to those who schedule ahead of time with me. Scheduling also allows me to divide meeting times so that I can provide ample individual time to each student.

**Dr. Waddell's Email policy:** Please feel free to email me with any and all questions you have. I will reply to emails within 24 hours (usually within an hour or so). I will not respond to any emails after 8pm, so if you email later at night, expect a reply first thing in the morning.

### Appendix 2 – Schedule

| A Virtual CURE | | | | | | |
|---|---|---|---|---|---|---|
| **Week/Module** | **Day** | **Date** | **Content Day Topic** | **Lab Meeting Day** | **JC article** | **Faculty Notes** |
| | | | | **No class** | | |
| **1** | W | 9/2 | Intro to Professors, students, and the course. | | | |
| **2** | T | 9/8 | What is Chronic Pain? A focus on the disease. | | | |
| | W | 9/9 | | What do scientists do? How do you become one? | | Introduce students to the academic process of becoming a scientist. Discuss career paths. |
| **3** | M | 9/14 | What is Addiction? A focus on the disease | | | |
| | W | 9/16 | | How do they read research articles??? | | Introduce students to navigating research articles. |
| **4** | M | 9/21 | What is the BioPsychoSocial Aspect of Human Health? | | | |
| | W | 9/23 | | How do scientists ask questions? Hypotheses, dependent variables, failure, etc. Also RCR and Lab Safety Training. | "Social transfer of pain in mice", Science Advances, 2016 | Introduce the concept of developing a research question, hypothesis, and prediction based on an observation of nature. Also discuss how to design experiments. |
| **5** | M | 9/28 | Basic Cell Biology - what is a cell? What is a neuron? | | | |
| | W | 9/30 | | Designing our study - asking the right question within our limits. | "Nerve injury drives a heightened state of vigilance and neuropathic sensitization in Drosophila", Science Advances, 2019 | Introduce students to our actual semester-long research project. |
| **6** | M | 10/5 | Basic Molecular Biology - what are signaling cascades? GPCRS? | | | |
| | W | 10/7 | | continuing discussing what we're doing for this. | | Continue discussing how the research project will work. |
| **7** | M | 10/12 | Basic Neurophysiology | | | |
| | W | 10/14 | | **Proposal Presentation** | "Drosophila melanogaster foraging regulates a nociceptive-like escape behavior through a developmentally plastic sensory circuit", PNAS, 2019 | Students should receive "lab in a box" this week in preparation of using them the following week. |
| **8** | M | 10/19 | Basic Neuroanatomy of the CNS | | | |
| | W | 10/21 | | Climbing demo, anesthetizing flies, flipping flies, sexing flies. | | Demonstrate basic fly pushing. Have students enter break out rooms to work together. |
| **9** | M | 10/26 | Basic Neuroanatomy of the PNS | | | |
| | W | 10/28 | | Sensitivity demo | | |
| **10** | M | 11/2 | Basic Neuropharmacology - what are neurotransmitters? | | | |
| | W | 11/4 | | **Research in Progress 1** | "Naltrexone Reverses Ethanol Preference and Protein Kinase C Activation in Drosophila melanogaster", Frontiers in Physiology, 2018 | |
| **11** | M | 11/9 | Basic Neuropharmacology - what are neurotransmitter receptors? | | | |
| | W | 11/11 | | Tolerance demo, Make food | | Consider having students flip flies to new tube to keep stocks healthy. |
| **12** | M | 11/16 | Basic Pain Neurobiology | | | |
| | W | 11/18 | | CAFE assay demo | | **This is the hardest assay to perform.* |
| **13** | M | 11/23 | Chronic Pain - Working with patients and our community | | | |
| | W | 11/25 | **Thanksgiving - No Class** | | | |
| **14** | M | 11/30 | Basic Reward Neurobiology | | | |
| | W | 12/2 | | **Research in Progress 2** | | |
| **15** | M | 12/7 | Addiction - Working with patients and our community | | | |
| | W | 12/9 | | **Thesis Presentation** | | |
| **Finals** | | | **Conference Debrief Presentation - Monday, Dec. 21, 2020 2:45pm-5:45pm** | | | |

| Pre-lecture videos | |
|---|---|
| **Module** | **Website** |
| 2 | What is pain? |
| | Would you opt for a life without pain? |
| | The mysterious science of pain |
| | How does your brain respond to pain? |
| | The mystery of Chronic Pain |
| 3 | Understanding the chemistry of the brain |
| | How do drugs get into the brain? |
| | How and WHY does addiction form? |
| | How do opioids affect the brain? |
| | Why is the opioid epidemic so hard to address? |
| | Addiction is a disease and needs to be treated as such. |
| 4 | What is the BioPsychoSocial Aspects of Human Health? |
| | What does the BPS aspects of human health look like in practice? |
| | How your zipcode defines your health |
| | How the BPS model affects pain |
| | How the BPS model affects addiction |
| | Should we just legalize all addictive substances!? |
| 5 | Cell Biology Introduction |
| | Parts of the Cell Overview |
| | Cell Membrane Introduction |
| | Anatomy of a Neuron |
| 6 | Signaling Cascades |
| | Membrane Receptors |
| | G-Coupled Protein Receptors |
| 7 | Resting Membrane Potential |
| | Action Potentials Part 1 |
| | Action Potentials Part 2 |
| | Sodium Potassium Pump |
| | Synaptic Structure |
| | Neurotransmitter Release |
| | Types of Neurotransmitters |
| | Neurotransmitter Receptor Types |
| 8 | Structure of the Nervous System Overview |
| | Functions of the Nervous System |
| | Central Nervous System Overview |
| 9 | Peripheral Nervous System Overview. |
| | Autonomic vs Somatic Nervous System Overview |
| | Overview of the Peripheral Nervous System |
| 10 | The Nervous System Part 1 |
| | The Nervous System Part 1 |
| | The Nervous System Part 1 |
| 11 | Pharmacology - Pharmacokinetics (PK). |
| | Pharmacology - PHARMACODYNAMICS (PD) |
| 12 | Nociceptors - An Introduction to Pain. |
| | PAIN! Physiology - The Ascending Pathway, Descending Pain Pathway and the Substantia Gelatinosa |
| 13 | Chronic Overlapping Pain Conditions: Definition and Causes |
| | The Psychology of Pain: For Patients and Practitioners |
| | The Grief of the Chronic Pain Patient |
| | Cannabinoids and Current Data |
| | Central Sensitization and Opioid Tapering |
| 14 | Opioid Drugs, Part 1: Mechanism of Action. |
| | Opioid Drugs, Part 2: Addiction and Overdose |
| 15 | Fudin v Gudin, Part 2: Cannabis, Kratom, and Opioid Guidelines. |
| | Opioid Use in Headache Management |
| | Beyond Opioids: A Plan B for Pain |
| | Educating About Opioids |
| | The Opioid Pendulum: Finding a Balance |
| | I was in opioid withdrawal for a month — here's what I learned \| Travis Rieder \| TEDxMidAtlantic |

### APPENDIX 3 – JOURNAL CLUB WORKSHEET

#### Journal Club Worksheet

Instruction: Give this week's assigned paper a read and answer the following questions. Since many of you may not have read a research article before, this will be challenging. Even the most experienced researchers have trouble reading research articles outside their immediate field. Instead of getting stuck on the minutia, try your best to have a clear understanding of the overall paper and the general goals, findings, and details of each figure.

1. What is the title of this article?
2. What is the name of the scientific journal this article is published in?
3. What university is the first author from?
4. Write a summary of the article in your own words in approximately 100-300 words. Your summary should include the experimental question, main hypothesis, methodology, results, and conclusions. Try to distill out the logic of the article's study.
5. Write down two questions you'd like to ask the presenting group.

### APPENDIX 4 – LAB MANUAL

### Lab Manual: This manual contains all necessary components for carrying out this semester-long research project.

**Project Description:**

Chronic pain is a huge public health concern. Those suffering from chronic pain are more at risk of developing a substance use disorder, but we don't know what biological, psychological, or social aspects are involved in increasing this risk. Is it that the chronic pain causes patients to seek out relief at a higher rate or is it that neurological circuits involved in both chronic pain and reward overlap and become rewired? To explore this, we need an animal model. Drosophila experience both pain and addiction so they are a reliable animal to model this relationship behaviorally. However, this relationship has never been tested in Drosophila before. So that's what we're going to do- literally for the first time EVER.

Our **observation** is that pain and addiction tap into similar brain areas involved in learning, memory, reward, and placing affect on different external experiences.

Our **research question** is whether pain makes addiction more likely to occur because it interacts with these similar neuronal structures.

Our **hypothesis** is that Drosophila experiencing pain will have a higher preference for rewarding substances (in this case, the alcohol in hand sanitizer) while Drosophila that can't experience pain will have a reduced preference.

We'll test this in four separate groups of Drosophila:

1. **WT**: wild-type flies
2. **PL**: Painless flies that can't feel pain because they've been genetically engineered to lack the neurons that transmit pain
3. **WT + A**: wild-type flies that have undergone a leg amputation to induce chronic pain
4. **PL + A:** Painless flies that have undergone a leg amputation to induce chronic pain.

If our hypothesis is supported, we **predict** that WT animals that have experienced a leg amputation to induce chronic pain will feature greater "addiction-like behaviors" than WT animals that are not experiencing pain. We would also **predict** that PL animals that have experienced amputation but no pain would feature similar "addiction-like behaviors" to WT animals that have no pain.

To test for these "addiction-like behaviors", we will use four separate behavioral assays that assess Drosophila's preference for alcohol. Although alcohol is not the same as opioids, it taps into similar reward circuitry as opioids and can thus serve as a safe way for us to explore addiction generally:

1. **Negative Geotaxis Assay**: Because all our behavioral assays require the flies climb to the top of the tube, we need to know if their mobility or ability to do this is affected by a leg amputation. If it does affect their mobility, then the amputation could interfere with our results and is thus not a valid method to run our behavioral assays. Download the Negative Geotaxis Video from Github for visual guidance.
    a. **Paper**: Gargano, J. W., Martin, I., Bhandari, P. & Grotewiel, M. S. Rapid iterative negative geotaxis (RING): a new method for assessing age-related locomotor decline in Drosophila. Exp Gerontol 40, 386–395 (2005).

2. **Sensitivity Assay**: This will test whether there are any differences in sensitivity to EtOH. Studies demonstrate that lower sensitivity to alcohol increases the chances an organism will develop alcohol-use disorder. Download the Sensitivity and Tolerance Assay Video from Github for visual guidance.
    a. **Paper**: Bhandari, P., Kendler, K. S., Bettinger, J. C., Davies, A. G. & Grotewiel, M. An assay for evoked locomotor behavior in Drosophila reveals a role for integrins in ethanol sensitivity and rapid ethanol tolerance. Alcohol Clin Exp Res 33, 1794–1805 (2009).
3. **Tolerance Assay**: This will test how quickly drosophila develop tolerance to EtOH. Studies demonstrate that increased development of tolerance can increase the chances an organism will develop alcohol-use disorder. Download the Sensitivity and Tolerance Assay Video for visual guidance.
    a. **Paper**: Bhandari, P., Kendler, K. S., Bettinger, J. C., Davies, A. G. & Grotewiel, M. An assay for evoked locomotor behavior in Drosophila reveals a role for integrins in ethanol sensitivity and rapid ethanol tolerance. Alcohol Clin Exp Res 33, 1794–1805 (2009).
4. **CAFÉ Assay**: This assay will test Drosophila preference for EtOH over sugar water. Studies demonstrate that an increased preference of EtOH is a sign of alcohol-use disorder. Download the CAFÉ Assay video from Github for visual guidance.
    a. **Paper**: Devineni, A. V. & Heberlein, U. Preferential Ethanol Consumption in Drosophila Models Features of Addiction. Curr Biol 19, 2126–2132 (2009).

To take care of these flies and set up the experiments, we also need to become familiar with a few standard protocols:

1. **Fly Pushing**: Handling flies is called "fly pushing". You need to be comfortable transferring flies from one container to another, sexing flies, and other protocols. A fantastic resource to do this was published in the journal JUNE a few years back. Download the Anesthetize Fly Video from Github for visual guidance.
    a. **Paper**: Pulver, S. R. & Berni, J. The Fundamentals of Flying: Simple and Inexpensive Strategies for Employing Drosophila Genetics in Neuroscience Teaching Laboratories. J Undergrad Neurosci Educ 11, A139–A148 (2012).
2. **Leg Amputation**: A validated method of inducing long-term chronic pain in Drosophila is to amputate one leg. This can be done carefully with a sewing needle.
    a. **Paper**: Khuong, T. M. et al. Nerve injury drives a heightened state of vigilance and neuropathic sensitization in Drosophila. Science Advances 5, eaaw4099 (2019).

**Finding Papers**: The University has subscriptions to most all research journals. When on campus, you can freely access them via the internet you connect to. When at home, however, you're out of this network. To gain access to papers, you'll have to log in through the library to gain access to the papers for free.

### Lab in a Box

For this semester-long study, you will each receive a lab in a box that includes all necessary components to carry out this entire semester-long experiment. If any of your components leak or you need replacement pieces, just ask.

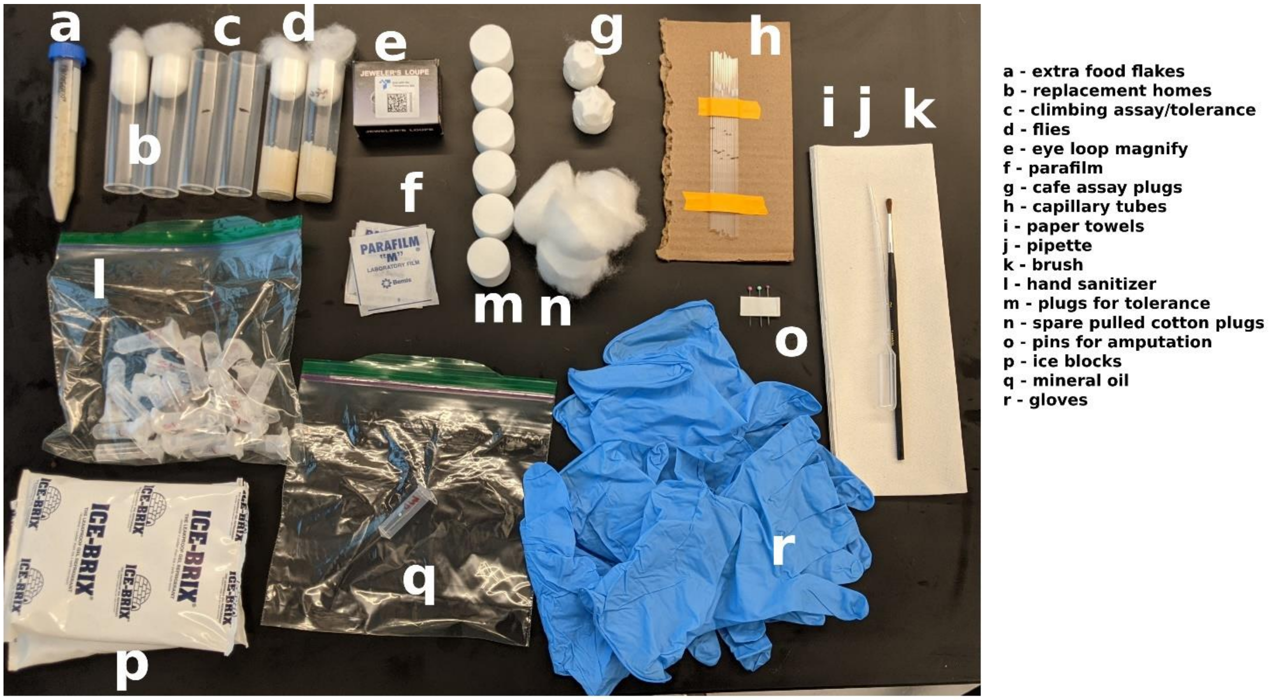

A- Extra food flakes: These flakes are dry food. Just add water in a 1:1 ratio to prep food for the flies.

B- Replacement Homes: These are replacement vials for when you need to transfer flies to new homes with fresh food.

C- Climbing Assay/Tolerance Chambers: These are clean vials for you to perform these assays. They are pre-marked at the 5 cm location for the climbing assays.

D- Flies: These are you fruit flies. You will receive two genotypes: Wild-Type and Painless. They already have food in them.

E – Eye loop Magnifiier. This is an eye loop that watchmakers and jewelers use to work on tiny objects. They're perfect for doing work with Drosophila.

F- Parafilm: This is similar to plastic wrap, but will help you ensure a tight seal over tubes so that nothing leaks and nothing evaporates.

G- Café Assay Flugs: These "fly plugs" aka Flugs will be used to soak your EtOH and place on the top of your behavioral tubes so that the flies can be exposed to the ethanol vapors. These tubes have had holes poked into them so you can easily slip the capillary tubes into them.

H- Capillary tubes: For the CAFÉ assay, you'll need to feed the flies. The glass capillary tubes will allow you to measure the amount of food they eat.

I- Paper Towels: Just paper towels to keep things clean and keep your flies dry.

J- Pipette: You can use this pipette to transfer solutions.

K- Brush: Once flies are anesthetized, you can use this paint brush to move them around.

L- Hand sanitizer and Sucrose: These tubes contain a range of 0-50% ethanol and 5% sucrose for your experiments. Each tube is labelled with what is inside of it.

M- Flugs for Tolerenace: These flugs are used in your tolerance assay. They don't have any holes poked into them.

N- Spare Pulled Cotton Plugs: These pulled cotton plugs can be used to close the fly home vials. They allow air to pass, but not the flies.

O- Pins for Amputation: These sewing pins can be used to amputate the legs of flies for inducing chronic pain.

P- Ice blocks: To anesthetize flies, you'll put them on ice.

Q- Mineral oil: To load the capillary tubes with your solutions, you'll need to pre-wet the tip with mineral oil.

R- Gloves: While the experiments do not contain any hazardous materials, we provided gloves in case you'd like to stay clean.

### Protocol: Anesthetizing Flies for Fly Pushing

**Purpose:** To move flies from one tube to another, you must first knock them out. Uniquely, when flies experience cold temperatures, they fall asleep. We have a video attached to show you the procedure as well.

**Materials Needed**:

*Drosophila* in their home tube.

Ice pack

Paper towel

Paint brush

**Procedure**:

1. Take ice pack out of the freezer and wrap with paper towel. Drosophila will be placed directly on this ice pack and so the paper towel will keep them dry.
2. In one hand, tap the Drosophila in their home tube down so that the flies fall to the surface of the food.
3. Quickly remove the cotton plug and flip the tube completely over so that no flies escape.
4. Tap the side of the tube to ensure all flies have fallen to the ice pack. Be careful not to let the flies escape.
5. Watch the flies as they begin to stop moving. They will fall asleep within 20-30 seconds and will stay asleep as long as they are on the ice.
6. Remove the tube and use the paint brush to move them around to where they need to be. Flies can stay on ice for up to 10-15 minute before dying so don’t keep them there too long!

### Protocol: Leg Amputation

**Purpose:** To induce a state of chronic pain in the animals, we must induce an injury. We can do this while reducing the chance of mobility issues by amputating the right middle leg of the fly at the femur segment. This has been shown to induce a state of chronic pain and hypervigilance starting at day 7 following the amputation and lasting up to 21 days later.

**Materials Needed**:

*Drosophila* in their home vial.

Ice pack

Paper towel

Paint brush

Sewing needles

Eye loop

**Procedure**:

1. Anesthetize flies on ice.
2. Use the paint brush to position flies on their backs.
3. Amputate the femur segment of the right middle leg with two sewing needles by crossing the needles and pressing down on their leg. The Drosophila leg is made up of three segments. The femur is the closest segment to the body. This is not an easy process so don't get too frustrated as you're doing it. You may need to use the eye loop to see the flies more closely. These eye loops can be held in your eye socket like a watchmaker or held up with a piece of tape or other apparatus in a position that allows you to look at the flies.

### Behavioral Assay: Negative Geotaxis Assay

**Purpose:** Drosophila have a tendency to run away from the force of gravity. This is called negative geotaxis and is seen as the flies move up the side of the tube after being hit down to the bottom. This behavior is essential for all our behavioral assays so we need to confirm the flies are able to do this after injury. A video is included to demonstrate this process

**Materials Needed**:

*Drosophila* in their home vial.

Clean vial pre-marked to 5 cm.

Pulled cotton plug

Timer

**Procedure**:

1. Flip the flies from their home vial into a clean vial pre-marked at 5cm from the bottom. This avoid anesthetizing flies while transferring them from one vial to another.
    a. To flip the flies, tap the flies to the bottom of their home vial and remove the pulled cotton plug.
    b. Quickly take the clean pre-marked vial and place it on top so the opening on both meet.
    c. Flip the entire system over carefully to not let any flies lose.
    d. Tap the flies into the clean vial and put the new pulled cotton plug onto the clean vial to keep flies inside.
2. You can now begin your experiment. Tap the flies down to the bottom of the tube and set the timer to 18 seconds.
3. Count the number of flies that move across the 5 cm mark and use this number to calculate the percentage of flies that were negative geotaxic.
4. Repeat steps 2-3 for a total of 3 times. Your calculation will be the average of your three trials.

### Behavioral Assay: Sensitivity Assay

**Purpose:** To perform this assay, you will expose the Drosophila to increasing levels of ethanol that are administered via vapor by soaking the ethanol into the cotton flug. As the flies experience higher concentrations of EtOH, they will begin to fail the negative geotaxis assay. Flies that are more sensitive to EtOH, will fail sooner than those that are not as sensitive. A video is included to demonstrate this process.

Note: This assay will take approximately 2 hours per genotype. Also, if you have two timers, it will make this easier for you. Set one timer to 1 minute and the second timer to 18 seconds. If you only have one timer, you will have to switch back and forth.

**Materials Needed**:

Timer (2)

Empty premarked vials (2) with cotton plugs (3)

Pulled cotton plugs (1)

Both *Drosophila* genotypes

Eppendorf tubes containing: 0% Ethanol, 25% Ethanol, and 50% Ethanol (2 each)

Notebook

**Procedure**:

1. Starting with one of the genotypes, gather an empty premarked vial, a 3 hard cotton plugs, one pulled cotton plug, both timers, and one Eppendorf tube for 0%, 25% and 50% Ethanol.
    a. Start with 0% Ethanol. On a clean cotton plug, dump the contents of the Eppendorf tube onto one side. (Note: That will be the side that you put into the vial.)
        i. Label the top of this cotton plug with the Ethanol concentration that was used on it (you will reuse this for the experiment with the second genotype).
    b. Remove 5-10 flies from your genotype stock (depending on how many flies you have available).
    c. Place those flies into the clean vial and cap with the pulled cotton plug to allow them to wake up (not the plug with the Ethanol content on it).
    d. Immediately place all of the remaining flies back into their original vial with food. (Note: Do not leave the flies on the ice for longer than 2 minutes if possible).
2. Tap the flies down in the empty premarked vial and replace the pulled cotton plug with the saturated cotton plug (again, make sure the wet end goes into the vial).
3. You will now perform the negative geotaxis assay with these flies (refer to negative geotaxis protocol).
4. Following the negative geotaxis assay trial, allow the flies to rest in the vial for one minute.
5. After a 1-minute rest period, repeat the negative geotaxis assay.
6. Repeat step 5, 18 more times. This will give you 20 trials (20, 1-minute time points) for this genotype at this specific ethanol concentration.
7. Repeat steps 2-6 for the same genotype using 25% and 50% ethanol. Do not reuse the same flies from any of the previous trials.

   a. You should have two premarked empty vials. Switch back and forth between them during this experiment.
   b. Leave the flies you used in the first part of this assay in the empty premarked vial they are in for now.
   c. Remove 5-10 new flies from your genotype stock (depending on how many flies you have available).
   d. Place those flies into the second empty premarked clean vial and cap with the pulled cotton plug to allow them to wake up.
   e. Immediately place all of the remaining flies back into their original vial with food. (Note: Do not leave the flies on the ice for longer than 2 minutes if possible).
   f. Now dump the flies in the first empty premarked vial onto the ice pad. Move them to a corner of the pad. These flies can remain on the ice pad for the remainder of this experiment. (Note: You do not want to reuse flies between Ethanol concentration groups, therefore do not mix these flies back into the stock yet.
   g. Wipe out the now unused empty premarked vial prior to using again.
8. When you are done with all three Ethanol concentration for this genotype, place all flies that were used back into the genotype stock. Let the vial rest on its side until all flies have awoken from the ice.

End experiment for genotype 1.

Now, repeat steps 1-8 with the second genotype. Reuse the saturated cotton plugs from the first experiment for this experiment.

### Behavioral Assay: Tolerance Assay

**Purpose:** To perform this assay, you will expose Drosophila to a high concentration of ethanol multiple times. Those that have developed a tolerance to it will recover faster and be able to perform the negative geotaxis assay sooner than those that did not become tolerant to the EtOH. This assay includes the same video as seen in the sensitivity assay, but changes the way the animals are exposed to EtOH.

Note: This assay will take approximately 5 hours per genotype. You will have a four-hour rest period in the middle of this assay.

**Materials Needed**:

Timer (2)

Empty premarked vials (2) with cotton plugs (3)

Pulled cotton plugs (1)

Both *Drosophila* genotypes

Eppendorf tubes containing: 0% Ethanol and 50% Ethanol (4 each)

Notebook

**Procedure**:

1. Starting with one of the genotypes, gather an empty premarked vial, 2 hard cotton plugs (the ones used during the first sensitivity assay), one pulled cotton plug, both timers, and one Eppendorf tube for 0% and 50% Ethanol.
    a. Start with 0% Ethanol. On the cotton plug, dump the contents of the Eppendorf tube onto one side. (Note: That will be the side that you put into the vial.)
    b. Remove 5-10 flies from your genotype stock (depending on how many flies you have available).
    c. Place those flies into the clean vial and cap with the pulled cotton plug to allow them to wake up (not the plug with the Ethanol content on it).
    d. Immediately place all of the remaining flies back into their original vial with food. (Note: Do not leave the flies on the ice for longer than 2 minutes if possible).
2. Tap the flies down in the empty premarked vial and replace the pulled cotton plug with the saturated cotton plug (again, make sure the wet end goes into the vial).
3. You will now perform the negative geotaxis assay with these flies (refer to negative geotaxis protocol).
4. Following the negative geotaxis assay trial, allow the flies to rest in the vial for one minute.
5. After a 1-minute rest period, repeat the negative geotaxis assay.
6. Repeat step 5, 28 more times. This will give you 30 trials (30, 1-minute time points) for this genotype at this specific ethanol concentration.
7. Tap the flies down and replace the ethanol saturated cap with a pulled cotton cap. Allow the flies to rest in the vial for four hours.
8. During the four-hour rest, repeat steps 2-6 for the same genotype using 50% ethanol. Do not reuse the same flies from the previous trial.
    a. You should have two premarked empty vials.

   b. Remove 5-10 new flies from your genotype stock (depending on how many flies you have available).
   c. Place those flies into the second empty premarked clean vial and cap with the pulled cotton plug to allow them to wake up.
   d. Immediately place all of the remaining flies back into their original vial with food. (Note: Do not leave the flies on the ice for longer than 2 minutes if possible).
9. After a four-hour rest period, resaturate the hard cotton plug with 0% ethanol concentration. Tap the flies in the 0% ethanol vial down and replace the pulled cotton plug with the newly saturated hard cotton plug.
10. Repeat the sensitivity assay on these flies again until all of the flies are knocked out.
11. Repeat steps 9-10 for the 50% ethanol concentration trial.
12. When you are done with both Ethanol concentrations for this genotype, place all flies that were used back into the genotype stock. Let the vial rest on its side until all flies have awoken from the ice.

End experiment for genotype 1.

Now repeat steps 1-12 with the second genotype. Reuse the saturated cotton plugs from the first experiment for this experiment.

### Behavioral Assay: CAFÉ Assay

**Purpose:** This assay will test whether the flies have a preference for EtOH over sugar water. Flies that have developed addiction-like behaviors will have a preference for EtOH over the sugar water. We can measure this by placing the flies into a tube that has no food. We can then insert capillary tubes filled with either ethanol + sucrose or sucrose alone. After leaving the flies in these tubes overnight, they will choose which tube to drink from. The following day, you can measure the amount of liquid that was consumed from each capillary tube by measuring how far the meniscus has dropped. If the flies preferred the ethanol-dosed food more, then the meniscus would have dropped lower than that in the capillary tube with sucrose only.

Note: This experiment is broken into two parts over the course of two days. First, we will discuss the experimental setup. Second, we will perform the experiment and calculate our data.

**Materials Needed**:

Timer

Empty vials (2)

Shaved hard cotton plugs

Microcapillary Pipets (4)

Eppendorf Tubes (15% Ethanol + sucrose and 5% sucrose) (2 each)

Both *Drosophila* genotypes

Notebook

**Procedure**:

Part 1:

1. Take both shave hard cotton plugs with the holes poked through the top of them and **GENTLY** insert a microcapillary pipet through each hole (4 holes, 4 pipets, 2 per cotton plug).
    a. Make sure that the white mark on the microcapillary tube is sticking out of the top of the vial (the none shaved side).
2. Mark one microcapillary pipet per cotton plug (this will allow you to differentiate which pipet contains the 15% ethanol concentration).
3. One cotton plug at a time, fill each microcapillary tube (one with 5% sucrose and one with 15% ethanol + 5% sucrose) using capillary action.
    a. To fill each pipet, place the bottom of the pipet into the Eppendorf tube and allow the pipet to draw in the liquid using capillary action. This may take a minute to completely fill.
    b. Mark the level at which the liquid is in the pipet. This should be slightly below the white mark on the capillary pipet.
    c. Once the liquid is in the pipet, do not allow the bottom edge to touch a surface. This will draw the liquid back out of the pipet.
4. Once both pipets are filled in a cotton plug, place it directly into an empty vial. (Don't press it down hard into the vial yet.)
    a. Fill the pipets for both cotton plugs at this time.

5. Each vial will house one genotype of our flies.
6. Knock out your flies using the ice pads and place an even number of flies (~20 flies) into each vial.
    a. If you only have 5-10 flies in one genotype, then only use the same number for the other genotype. Make sure that the number of flies in each empty vial are even.
7. Place those flies into an empty vial and firmly secure the cotton plug with the microcapillary pipets into the vial.
    a. Make sure you label each vial in some way to enable you to know which vial contains which genotype.
    b. Any flies not placed in the empty vials, immediately place back into their stock vials with food.

Part 2:

1. Allow the flies to wake up from the ice completely. This may take up to 20 minutes.
2. Once all of the flies have begun to move around again, you will start a timer for 24 hours.
3. After each hour, you will mark the liquid line in each microcapillary pipet.
    a. After a little bit, you may have to pull the pipet up slightly in order to see the liquid line. This is fine to do, just make sure the end of the pipet does not come into contact with the side of the vial or the cotton plug.
    b. If the liquid line is too low to pull the pipet up safely, then **GENTLY** tap the flies down in the vial and replace the shaved cotton plug with a pulled cotton plug. Push the pipet down through the cotton plug in order to see the liquid line, mark it, and then quickly replace the shaved cotton plug back into the vial.
4. After 24 hours, each microcapillary pipet should have 25 marks on it (Time Point 0-24).
5. Remove the flies from each empty vial using the ice packs and place them back into their stock vials.
6. Remove the microcapillary pipets from the shaved cotton plugs.
    a. Make sure you know which pipets came from which genotype.
7. Calculate your data.
    a. The length of each pipet's liquid reservoir (base to middle of the white area) is 9 cm. The total volume that can be held by each pipet when that reservoir is completely filled is 5 ul. Using that, calculate the volume of liquid at each time point for each pipet.
8. Record your data for this experiment.

### Appendix 5 – Lab in a Box Assembly Plan

| Stock items for purchase | | | | | | | | |
|---|---|---|---|---|---|---|---|---|
| **Item** | **Company** | **Item Number** | **Unit** | **URL** | **Price (2021)** | **Quantity** | **Total** | **Notes** |
| VWR® Drosophila Vial, Wide | VWR | 75813-156 | 500 | https: | $ 77.78 | 1 | $ 77.78 | |
| VWR® Drosophila Vial Plugs, Cellulose Acetate | VWR | 89168-888 | 1000 | https: | $ 129.85 | 1 | $ 129.85 | |
| Cotton Balls - Medium | VWR | 10030-842 | 4000 | https: | $ 56.87 | 1 | $ 56.87 | |
| 15 ml falcon tubes | VWR | 62406-200 | case of 500, 50/bag | https: | $ 303.56 | 1 | $ 303.56 | Any tube is fine. This is just for transporting fly food. |
| KINGMAS Pocket Jewelry Loupe 30x 21mm Jewelers Eye Magnifying Glass Magnifier | Amazon | na | 1 | https: | $ 4.99 | 20 | $ 99.80 | Any eye loop is fine. Each student needs one. |
| Parafilm M, Bemis | VWR | 52858-076 | 1 | https: | $ 86.36 | 1 | $ 86.36 | |
| Ward's® Instant Drosophila Medium | VWR | 470024-740 | pack of 4 | https: | $ 24.99 | 1 | $ 24.99 | |
| VWR® Microhematocrit Capillary Tubes Heparin Coated | VWR | 15401-560 | 100 | https: | $ 36.09 | 2 | $ 72.18 | |
| Round Paint Brushes | Amazon | na | 50 | https: | $ 14.80 | 1 | $ 14.80 | |
| sewing pins | Amazon | na | 350 | https: | $ 5.99 | 1 | $ 5.99 | |
| paper towels | na | na | na | na | na | na | na | any paper towels are fine |
| gloves | na | na | na | na | na | na | na | any lab gloves are fine |
| ziploc bags | na | na | na | na | na | na | na | any bag to consolidate vials is fine. |
| mineral oil | Amazon | na | 4 oz | https: | $ 6.99 | 1 | $ 6.99 | any mineral oil is fine. Used for pre-wetting capillary tube tip. |
| Ice Pack | Amazon | na | 48 | https: | $ 21.67 | 1 | $ 21.67 | |
| ethanol | VWR | 97064-768 | 500 ml | https: | $ 51.70 | 1 | $ 51.70 | |
| Sucrose | VWR | BDH9308-500G | 500 g | https: | $ 88.54 | 1 | $ 88.54 | |
| Disposable transfer pipettes | VWR | 16001-170 | 500 | https: | $ 53.54 | 1 | $ 53.54 | |
| VWR® Microcentrifuge Tubes, Polypropylene | VWR | 87004-262 | 1000 | https: | $ 99.59 | 1 | $ 99.59 | |
| Painless Flies | Bloomington | 27895 | 1 | na | $ 15.00 | 1 | $ 15.00 | It is suggested to purchase flies a month in advance to expand the colony before shipping out. |
| Bloomington | Bloomington | 2376 | 1 | na | $ 15.00 | 1 | $ 15.00 | any Wildtype flies are fine. These are Oregon-R. It is suggested to purchase flies a month in advance to expand the colony before shipping out. |
| | | | | | | **Total** | **$ 1,224.21** | |
| **This is for a class of 20 but given the minimum unit size for some of these items, these should last for several years. The estimated cost per lab in a box is $15.* | | | | | | | | |

| Working items for each lab in a box | | |
|---|---|---|
| **Lab in a box item** | **quantity** | **notes** |
| A | 1 | Add 7 ml by volume of fly food flakes so that students can add 7 ml of water when needed to produce additional fly food |
| B | 2 | two extra tubes with pulled cotton tops |
| C | 2 | two behavioral tubes pre-marked at 5 cm for assays |
| D | 2 | Vials of WildType and vial of Painless flies. Don't mix genotypes. Ensure at least 20 flies are in tubes for reproductive success. |
| E | 1 | |
| F | 4 | 4 small sheets to reseal the ethanol is fine. |
| G | 2 | Should poke two holes for easy insertion of CAFÉ assay capillary tubes |
| H | 10 | tape capillary tubes onto a hard surface so they don't break |
| I | 3 | |
| J | 1 | |
| K | 1 | |
| L | 12 | For all experiments, each box will need 4 tubes of 0% EtOH, 2 tubes of 25% EtOH, 4 tubes of 50% EtOH, 1 tube of 15% EtOH + 5% Sucrose, and 1 tube of 5% Sucrose. |
| M | 6 | |
| N | 4 | |
| O | 3 | tape these needles together for safety and not to loose them. |
| P | 1 | |
| Q | 1 | |
| R | 3 | teach students to reuse gloves. |

### APPENDIX 6 – PEER EVALUATIONS QUESTIONS

**Peer Review of Presentations**

Instructions: Fill out the below review for the group's presentation on a rating of 1-10 with 1 being the worst and 10 being the best. 10 equates to a "100%" for that portion, 9 equates to a "90%" for that portion, etc etc.

Questions:

1. What presentation is this for?
2. What is your last name? (it will not be shared with the presenters. It's just for us to keep track of who's responding)
3. What is your first name? (it will not be shared with the presenters. It's just for us to keep track of who's responding)
4. Introduction/Background - did the presenters describe enough background for you to understand? (1-10 scale)
5. Results - did the presenters describe the results clearly for you to understand? (1-10 scale)
6. Conclusions - did the presenters clearly describe the conclusions of the paper sufficiently for you to understand it? (1-10 scale)
7. Questions - did the presenters address questions sufficiently? (1-10 scale)
8. Design - did the presenters design slides and communicate their messages well? (1-10 scale)
9. Goal – did the presenters accomplish the goal of the presentation as per the syllabus (1-10 scale)
10. What is one good thing the presenters did during their presentation? (open-ended)
11. What is one bad thing the presenters did during their presentation? (open-ended)

**Honors Neuro Peer/Self-Evaluation**

Instructions: Use this form to submit an evaluation of yourself as well as your peers. If you rate anyone at lower than a "3", it will trigger me to contact that person with your feedback.

Questions:

1. What is your name?
2. What would you rate your participation in this group so far? (5 = excellent, 4 = good, 3 = satisfactory, 2 = needs improvement, 1 = didn't participate)
3. Which group member are you evaluating?
4. What would the participation of this group member to the group assignment? (5 = excellent, 4 = good, 3 = satisfactory, 2 = needs improvement, 1 = didn't participate)
5. Would you like to leave any anonymous comments for this student?

*repeat questions 3-5 for multiple group members.

### APPENDIX 7 – RESEARCH PRESENTATION RUBRICS

**Proposal Presentation Faculty Rubric**

Presenter's Name(s):

Grader's Name (Optional):

Please assign an appropriate score for each category based on the following presentation.

In this presentation, students are expected to present a thorough background of the course's research project including the scientific rationale for the research question, hypothesis (or hypotheses), and all relevant background knowledge needed to understand the project. Additionally, students are expected to outline the research project including a timeline of experiments and what research question each experiment is designed to address.

Goal: Fully describe why and how the project is being pursued in such a way that stimulates interest in the work

| **Category** | **2** | **1** | **0** | **Score** |
|---|---|---|---|---|
| Preparedness | Student is completely prepared and has obviously rehearsed. | Student seems somewhat prepared, but lacking rehearsal. | Student does not seem prepared at all. | |
| Background Content | Presentation follows a logical flow connecting what is known in the field with outstanding questions. | Connections between what is known in the field and outstanding questions not established. | Background information is not supportive of outstanding questions in the field. | |
| Hypothesis | Hypothesis strongly structured based on literature support and cited data. | Hypothesis loosely structured based on literature support and cited data. | Hypothesis is not given or is stated but not supported by literature or data. | |
| Experimental Design | Each experiment is clearly laid out with a clear rationale and strong literature support. | Each experiment is minorly supported by the literature and experimental rationale is lmoderate. | Not all experiments are described and supported in full, or experimental layout and explanation are completely lacking. | |

| | | | |
|---|---|---|---|
| Proper Citations | All articles and background data are cited using a standardized citation method. | Some articles and background data are cited using a standardized citation method. | No citations present. |
| Ability to Answer Questions | Answered questions robustly with thoughts and connections beyond the original question. | Answered questions, but without depth or analysis. | Did not answer questions or answered questions incorrectly. |
| Ability to Defend Material Presented | Student able to strongly defend their presentation. | Student struggles to defend their presentation. | Student unable to defend their presentation. |
| Ability to Drive Class Discussion | Promoted robust discussion with class. | Promoted minimal discussion with class. | No class discussion promoted. |
| Presentation | Visual aids are used to enhance the presentation. All thoughts are articulated clearly. Student maintains audience interest. | Thoughts articulated clearly, however the presentation is not engaging. | Distracting, hard to understand. |

______/18

Additional Comments:

**Research in Progress Presentation Faculty Rubric**

Presenter's Name(s):

Grader's Name (Optional):

Please assign an appropriate score for each category based on the following presentation.

Students will present the class-aggregated results from the climbing and sensitivity assays. This will require collecting data from each class member's experiments, aggregating it, analyzing it, and making a conclusion based off it. Students will describe the experimental design, the scientific rationale for each specific experiment, and the results (in an appropriate graph or chart). Additionally, students should provide a brief reintroduction to the work including their scientific background, question and hypothesis (or hypotheses).

Goal: Provide an update on the first set of results and determine a conclusion for us to make decisions on the next steps of the project.

| **Category** | **2** | **1** | **0** | **Score** |
|---|---|---|---|---|
| Preparedness | Student is completely prepared and has obviously rehearsed. | Student seems somewhat prepared, but lacking rehearsal. | Student does not seem prepared at all. | |
| Background Content | Presentation follows a logical flow connecting what is known in the field with outstanding questions. | Connections between what is known in the field and outstanding questions not established. | Background information is not supportive of outstanding questions in the field. | |
| Hypothesis | Hypothesis strongly structured based on literature support and cited data. | Hypothesis loosely structured based on literature support and cited data. | Hypothesis is not given or is stated but not supported by literature or data. | |
| Experimental Design | Each experiment is clearly laid out with a clear rationale and strong literature support. | Each experiment is minorly supported by the literature and experimental rationale is moderate. | Not all experiments are described and supported in full, or experimental layout and explanation are completely lacking. | |

| | | | |
|---|---|---|---|
| Data Presentation | Data is presented well and discussed. Description of next steps for data collection are discussed. | Data is presented poorly and no conclusions can be drawn | Data is not presented. |
| Data conclusions | Data conclusions are supported by the data presented | Conclusions are made but are not supported by data | No concise conclusions are made |
| Ability to Answer Questions | Answered questions robustly with thoughts and connections beyond the original question. | Answered questions, but without depth or analysis. | Did not answer questions or answered questions incorrectly. |
| Ability to Drive Class Discussion | Promoted robust discussion with class. | Promoted minimal discussion with class. | No class discussion promoted. |
| Presentation | Visual aids are used to enhance the presentation. All thoughts are articulated clearly. Student maintains audience interest. | Thoughts articulated clearly, however the presentation is not engaging. | Distracting, hard to understand. |

______/18

Additional Comments:

**Thesis Defense Presentation Faculty Rubric**

Presenter's Name(s):

Grader's Name (Optional):

Please assign an appropriate score for each category based on the following presentation.

Students will create a final conclusion presentation about what these experiments conclude. Students are expected to present all data collected throughout the term. Students will describe the experimental design, the scientific rationale for each specific experiment, and the results (in an appropriate graph or chart). Additionally, students should provide a brief reintroduction to the work including their scientific question and hypothesis (or hypotheses). Finally, students will highlight the major conclusions of their work and potential future directions and implications of their work.

Goal: Describe the full picture of the project, it's conclusions, and future directions so we can assess where to go after this class is over.

| **Category** | **2** | **1** | **0** | **Score** |
|---|---|---|---|---|
| Preparedness | Student is completely prepared and has obviously rehearsed. | Student seems somewhat prepared, but lacking rehearsal. | Student does not seem prepared at all. | |
| Background Content | Presentation follows a logical flow connecting what is known in the field with outstanding questions. | Connections between what is known in the field and outstanding questions not established. | Background information is not supportive of outstanding questions in the field. | |
| Hypothesis | Hypothesis strongly structured based on literature support and cited data. | Hypothesis loosely structured based on literature support and cited data. | Hypothesis is not given or is stated but not supported by literature or data. | |
| Experimental Design | Each experiment is clearly laid out with a clear rationale and strong literature support. | Each experiment is minorly supported by the literature and experimental rationale is moderate. | Not all experiments are described and supported in full, or experimental layout and explanation are completely lacking. | |

| | | | |
|---|---|---|---|
| Data Presentation & Conclusions | Data is presented well and discussed. Description of next steps for data collection are discussed. Conclusions are supported by the data. | Data is presented poorly and no conclusions can be drawn | Data is not presented. |
| Final conclusions on the hypothesis | Conclusions are made to either refute or support hypothesis clearly. If the study is inconclusive, describe why. | Conclusions are made but are not supported by data and a complete picture is not made. | No concise conclusions are made |
| Bigger picture | Bigger picture is discussed. What does this mean for humans? How can this improve how we approach pain and addiction? What further studies could be conducted? | Bigger pictures is only touched on a little bit. | No bigger picture. |
| Ability to Answer Questions | Answered questions robustly with thoughts and connections beyond the original question. | Answered questions, but without depth or analysis. | Did not answer questions or answered questions incorrectly. |
| Presentation | Visual aids are used to enhance the presentation. All thoughts are articulated clearly. Student maintains audience interest. | Thoughts articulated clearly, however the presentation is not engaging. | Distracting, hard to understand. |

______/18

Additional Comments:

### APPENDIX 8 – JOURNAL CLUB WORKSHEET EXAMPLE

**Journal Club Worksheet – Student Example for "Social transfer of pain in mice", Science Advances, 2016.**

Instruction: Give this week's assigned paper a read and answer the following questions. Since many of you may not have read a research article before, this will be challenging. Even the most experienced researchers have trouble reading research articles outside their immediate field. Instead of getting stuck on the minutia, try your best to have a clear understanding of the overall paper and the general goals, findings, and details of each figure.

1. What is the title of this article?

"Social transfer of pain in mice"

2. What is the name of the scientific journal this article is published in?

"Science Advances"

3. What university is the first author from?

"Department of Behavioral Neuroscience, Oregon Health and Science University"

4. Write a summary of the article in your own words in approximately 100-300 words. Your summary should include the experimental question, main hypothesis, methodology, results, and conclusions. Try to distill out the logic of the article's study.

"In the journal article "Social transfer of pain in mice," Smith and her colleagues seek to answer the research question of whether "primary" mice's hyperalgesic state affects "bystander" mice that are in close proximity but not harmed. The authors' hypothesis was that there is social transfer of pain among mice. To test the hypothesis, Smith and colleagues injected the back-left paws of some mice with complete Freund's adjuvant (CFA), which they said "is well known to induce long-lasting, localized inflammation and hyperalgesia" and induced alcohol or morphine withdrawal in other mice. To confirm that the morphine dose created physical dependence, they dosed mice with naloxone, which induced withdrawal, confirming physical dependence. To establish alcohol dependence, researchers included one bottle of water and one bottle of alcohol with increasing ethanol concentrations. They compared whether there were differences in mice that were not exposed to CFA, morphine or alcohol based on whether they were housed and tested in the same room or another room as the mice who were exposed to CFA or alcohol. Researchers assessed whether hyperalgesia could be transferred due to non-visual cues by placing bedding from mice who experienced hyperalgesia in with mice who were housed in a separate room. Mice that were injected with saline, but housed in the same room as mice that were injected with CFA or withdrawing from morphine or alcohol displayed similar hypersensitivity. Mice that were not in the same room as mice who experienced hyperalgesia, but were exposed to their bedding, began experiencing hyperalgesia. Overall, the results indicate that pain is transferred socially among mice. When researchers introduced bedding from mice who experienced hyperalgesia to the cages of mice who had been housed in a separate room, the mice's pain response suggested that hyperalgesia transfers through smell among mice. "

5. Write down two questions you'd like to ask the presenting group.

"Did you observe any limitations in the study? Or could you expand upon some that the authors mentioned?

What were the differences in the sex that you observed amongst social pain transfer in mice? What might this mean for specifically for females that suffer from chronic pain?"

### APPENDIX 9 – RESEARCH PRESENTATIONS

#### Background

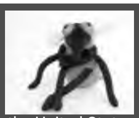

- "In 2016, about 20 percent of adults (50 million people) in the United States had chronic pain, defined as pain most days in the previous 6 months"
- "1 in 4 adults in chronic pain reports self-medicating with alcohol"
- In order to self medicate with alcohol, the patient would have to drink at a level equal to binge-drinking
- Opioids are usually used for physical pain, but patients tend to use it for emotional pain as well
- The current treatments for chronic pain are only 30% effective

#### What we Predict

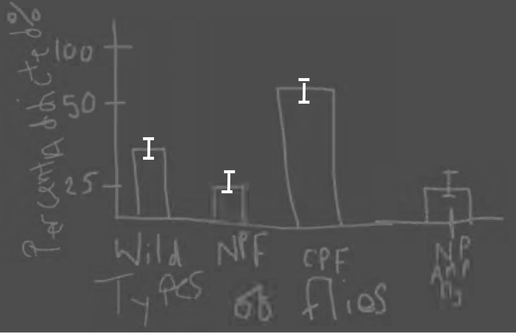

#### Protocol 2: Sensitivity/Tolerance Assay

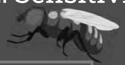

By tracking the sensitivity and tolerance to ethanol in the flies, we will be able to assess the genes involved in ethanol-related behavior.

1. eRING (ethanol Rapid Irritative Negative Geotaxis) is established as an assay for quantitating the sedative effects of ethanol on negative geotaxis.
2. Validated by assessing the acute sensitivity to ethanol and rapid ethanol tolerance in several different control strains and in flies with mutations
3. eRING also used in screen to ID mutants with altered ethanol-related behaviors.

Figure 1

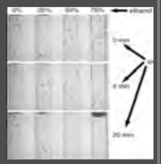

Prediction:

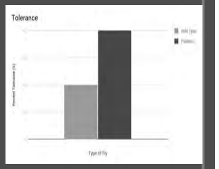

#### Why?

What led us to this study, Well we had observed that many people have sustained an injury while working or doing hobbies. This causes them to be prescribed opioids to relieve that pain. However, some people who get treated have chronic pain, and the opioids causes them to become dependent to feel okay. This causes the patients to seek out street drugs so that they can feel better.

Does having chronic pain increase the chance that one will become addicted to opiates and alcohol?

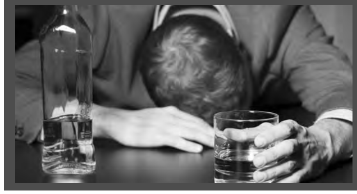

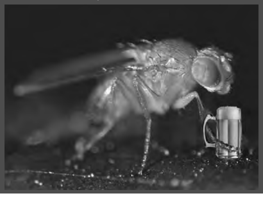

#### Hypothesis

- If you give alcohol to flies in chronic pain they will have a greater chance of developing an addiction.
- Null Hypothesis: If you give alcohol to flies in pain they will have the same chance of developing an addiction as flies not in pain.

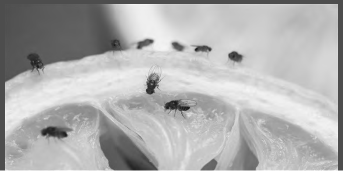

#### Protocol 1: Negative Geotaxis Assay

1.) Anesthetize the flies using $CO_2$ for 1-2 minutes and allow at least 18 hours for recovery after experiment
2.) Transfer the flies into each geotaxis tubes, splitting them into group A (wild type) and group B (painless flies) and place the tubes into the RING apparatus and secure with lid. Place camera 30 cm away from apparatus, a light should be behind apparatus with a filter to evenly distribute light in front of it (Figure A)
3.) After 1 minute, knock sharply on a table 3 times in succession to induce negative geotaxis responses. The position of the flies should be taken 4 seconds after initiating

By doing this negative geotaxis assay, we will be able to assess how far up the flies are able to travel up the tube, giving us a baseline of what the flies can do after triggering the escape response without being exposed to pain.

Prediction:

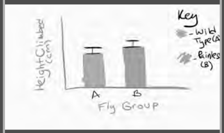

Figure A:

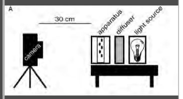

### Why The Fly?

#### Background Continued

Chronic pain is a huge public health concern. Those suffering from chronic pain are more at risk of developing a substance use disorder, but we don't know what biological, psychological, or social aspects are involved in increasing this risk. Is it that the chronic pain causes patients to seek out relief at a higher rate or is it that neurological circuits involved in both chronic pain and reward overlap and become rewired? To explore this, we need an animal model. Drosophila experience both pain and addiction so they are a reliable animal to model this relationships with behaviorally. This relationship has never been tested in Drosophila before.

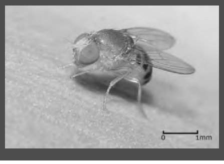

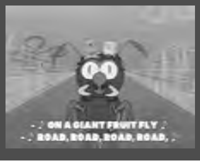

#### Protocol 0: Using Flies

Proper Care

it is important to keep fly stocks healthy. The simplest way to do this is to transfer flies to new vials every three weeks. This will promote breeding and inhibit growth of parasites. Secondly, keep careful track of humidity. Fruit flies prefer humid environments but food in vials tends to dry out and flies can slowly die of thirst. If fly food has lost its "creamy" consistency and/or pulled away from the vial itself, rehydrate immediately with a few drops of water.

Some fly lines are very unhealthy simply due to their mutations. Some mutants frequently get stuck in moist fly food. Adding dried baker's yeast and providing flies with emergent structures (bits of paper, Figure 2A) to climb and pupate away from food helps maintain sick lines.

The first step of any fly selection is anesthetization. In a dedicated fly facility, there are porous mats connected to a $CO_2$ source. When adults are put on the mat, they fall unconscious immediately. Flies can also be exposed to ether fumes or a commercial anaesthetic such as 'flynap; however, the simplest and least toxic (for flies and humans) alternative to $CO_2$ is cooling. This technique requires only ice, a freezer, petri dishes and paper. First, transfer flies to a new vial with or without food (cold-anesthetized flies may fall and stick to used, mushy food).

Then, transfer the vial to a freezer at −20°C. Wait until all flies fall to the bottom of the vial, but flies will start to die if held at −20°C for longer than 10 minutes. Meanwhile, fill a petri dish with ice, then cover with a second upside down petri dish and a piece of paper. This provides a cold-bottomed platform free of condensation. Place anesthetized flies on top of the paper and use a brush to gently select for appropriate genotypes. Individual flies will remain viable on this type of cold platform for 10–15 minutes.

### Drosophila Research Study: Data Presentation 1

### RECAP ON THE WORK

#### Scientific Background

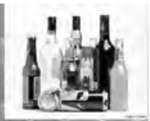

- 20% of adults report chronic pain in 2016
- Current chronic pain treatments are 30% effective
- Roughly 21-29% of people with chronic pain misuse opioids
- People self medicate with different drugs and alcohols
- In 2018 67,367 people died from drug overdose

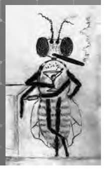

#### Protocol 4: Leg Amputation

- Right middle femoral amputation will be conducted to induce chronic pain in both the wild type strain of flies and the painless flies
- Amputation is known to result in neuropathic sensitization in the invertebrate, and fruit flies exhibit allodynia following peripheral nerve injury

Fruit flies, after having a leg amputated, respond viscerally to innocuous stimuli, and perceive stimuli that otherwise is not noxious as potentially detrimental

- *Drosophila melanogaster* display ethanol preference similar to alcohol addiction in humans, even when in an uninjured, healthy state
  - Amputation will be performed to see if this has any effect on the already pre-existing inclination towards alcohol that fruit flies possess

#### Leg Amputation Continued

We predict that the ethanol preference in *drosophila melanogaster*, paired with the peripheral nerve damage caused by femoral amputation, will result in amputated wild type flies (with chronic pain), group c, having an increased consumption of, and preference for, alcohol

- Alcohol dampens pain signals which sensory neurons send to the brain
  - Thus, flies with chronic pain may be more likely to become addicted due to the *incentive* provided by alcohol—numbing the chronic pain

We expect that amputation in painless flies will be less effective, in that the amputation will not greatly affect these flies since they cannot feel pain–thus the amputation will result in affected mobility, but likely not increase addiction

#### Prediction (Amputation)

We would expect the relative pain of amputated wild flies to be higher than pain of amputated painless flies. Relative pain could perhaps be measured based on responses to thermal nociception

- Based on group c having the highest level of pain, we would thereby expect them to have the greatest level of addiction

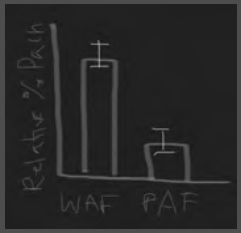

#### Protocol 3: CAFE Assay

In order to determine if the flies have developed an addiction to ethanol a CAFE assay will be tested. Each group of fly will be placed into vials with capillary feeders. Each capillary has sugar or ethanol (2 sugar, 2 ethanol). To setup the vial you will have 2-20 micropipette tips inserted into the top of the openings in the lid of the vial. Then fill 2 pipette tips with 5 microliters of ethanol and 2 with sugar water.

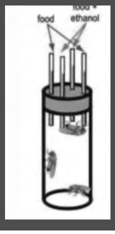

jove-121-55024-pmcvs_normal.mp4

What are we looking for?
When the flies are in the vials you will track the amount of time in minutes that your flies drink from either the sugar water or the ethanol. Along with tracking the amount of time spent at each feeder. You will also be tracking the amount of each substance is drunk. We are also looking to see after 5 days which feeder do the flies tend to drink from.

#### Prediction 1

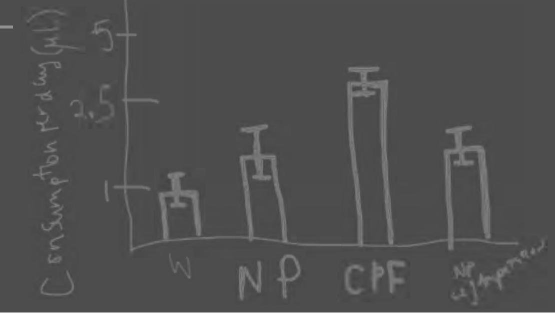

#### Prediction 2

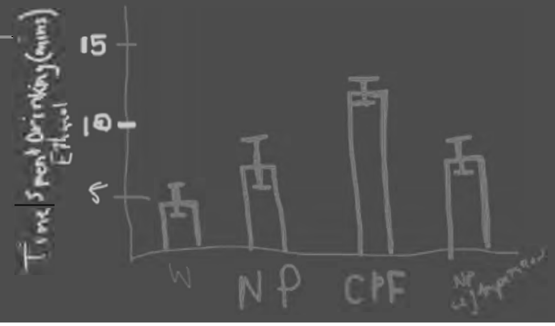

### Data for Negative Geotaxis Assay

#### Introduction to Results

- The goal of this assay is to see if there is a difference in the number of times flies from each group (wild type and painless) climb over the line after being tapped down, indicating a negative geotaxis escape response.
- With that, we can formulate the testable hypotheses:
  - Null hypothesis: Wild type negative geotaxis responses = Painless responses (i.e. there is no significant difference)
  - Alternative: Wild type escape responses ≠ Painless escape responses

9

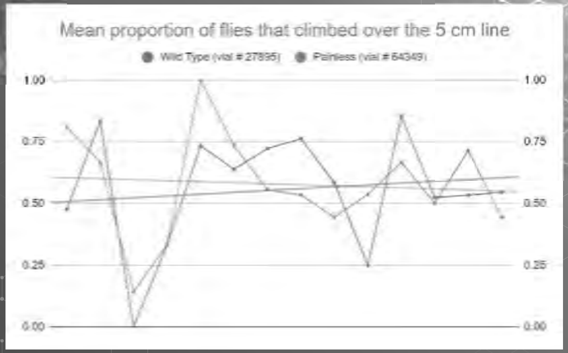

12

#### Hypotheses Made

- Previous hypothesis made: If you give alcohol to flies in chronic pain they will have a greater chance of developing an addiction.
- Null hypothesis: If you give alcohol to flies in pain they will have the same chance of developing an addiction as flies not in pain
- Additional hypothesis: If flies are constantly exposed to ethanol, eventually they will build up tolerance

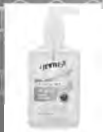

5

#### Experimental Design

1. Anesthetize the flies using CO2 or using the ice pack for 1-2 minutes
2. Transfer the flies into each geotaxis tubes, splitting them into two groups
   a. Group A (wild type)
   b. Group B (painless flies)
3. After 1 minute, knock sharply on a table 3 times in succession to induce negative geotaxis responses and turn on the timer
4. The position of the flies should be taken 18 seconds after initiating
5. Repeat the experiment for three trials per group of flies
6. Record your data on the excel sheet for Data Presentation 1

8

#### Conclusions

- Assuming an α of .05 (meaning a p-value < .05 is statistically significant), our p-value of 0.80679 is not significant, so we fail to reject the null hypothesis.
- We have no evidence to suggest the ability of *Drosophila* to feel pain affects the negative geotaxis escape response. The small difference between the means is likely caused by random variation
- In other words, there is no reason to think pain significantly affects the negative geotaxis response.

11

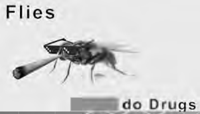

#### Question to be Studied

- We are studying if flies in chronic pain are more susceptible to becoming addicted to illicit substances.
- This study is important to research because if there is a link to chronic pain and an increase in drug use, people can find other ways to help out people that are in pain without jeopardizing their overall health.

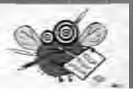

#### Scientific Rationale for Study

- The reason for this is to get a baseline understanding of how flies go up the tube
- For the next few experiments, the flies will be exposed to different things so we can compare the results of this assay to the next few assays

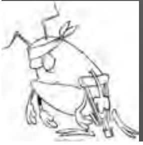

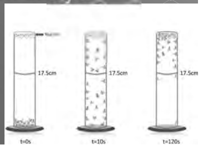

#### Results

| Wild type (vial # 27895) | Painless (vial # 64349) |
|---|---|
| Mean proportion of flies climbing above 5 cm line | Mean proportion of flies climbing above 5 cm line |
| 0.55621 (55.621%) | 0.57725 (57.725%) |

Difference of means
∓0.02104

p-value
0.80679

10

#### Scientific Rationale for Study

- The behavior changes due to alcohol are similar in both mammals and in fruit flies.
- Genes have been identified in fruit flies that affect the physiological responses to ethanol
- So this sensitivity assay will introduce ethanol to fruit flies and investigate the effects ethanol has on their negative geotaxis
- This will help better understand the effects ethanol has on locomotive responses

### Introduction to Tolerance

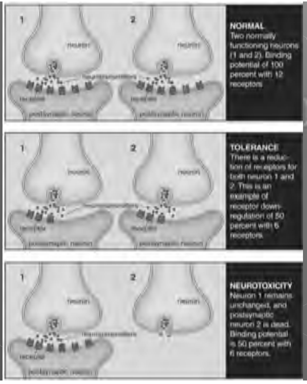

### Data for Sensitivity Assay

#### Data and Results (so far)

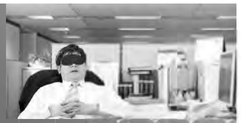

- No one has done the sensitivity assay yet

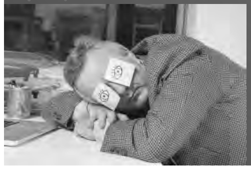

#### Tolerance in Action

- People who uses substances repeatedly build a natural immunity
- The natural immunity is a way to protect the body

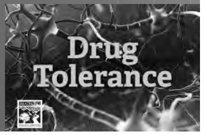

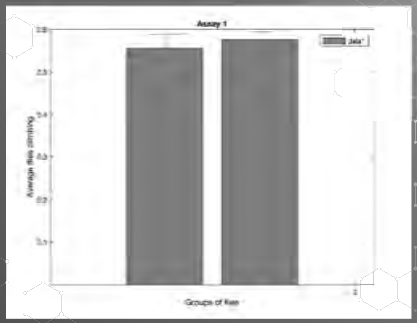

#### Experimental Design

1. Put the both groups of fruit flies asleep(painless and wild) by placing them on an ice pack for 1-2mins
2. Transfer the flies into tubes that have 5cm marked from the bottom
3. Cap the tube with a cotton plug doused in 0% (percentage of alcohol) hand sanitizer
4. Knock down the flies to the bottom and start an 18 sec timer. Record how many flies pass the 5cm mark after 18 sec
5. Repeat this process 20x with both fly groups
6. Then switch the hand sanitizer with 25% and 50% alcohol and repeat the same process 20x for each fly group
7. Record all 120 trials

#### Background on Tolerance

- According to the National Cancer Institute, tolerance is "A condition that occurs when the body gets used to a medicine so that either more medicine is needed or different medicine is needed."

#### The Experiment

- Uses the same method as the sensitivity assay
- Repetition of the Negative Geotaxis Assay will show tolerance develop.

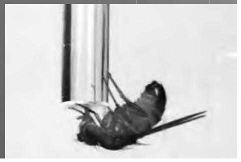

### Data Presentation 2

#### Opioids as treatment

Opioids are a class of drugs commonly used to treat chronic pain

- Relieves pain through blocking pain signals
- Can lead to opioid dependence and addiction
  - Has led to the Opioid Epidemic

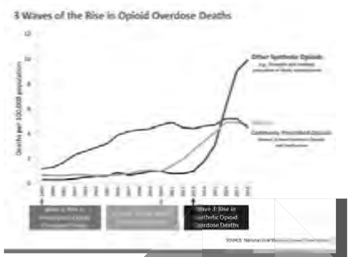

#### Drosophila and Alcohol Tolerance

- Flies are sensitive to alcohol
- Prolonged exposure should raise tolerance

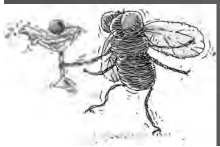

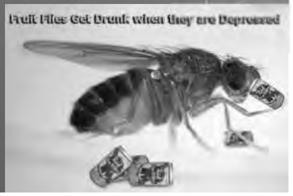

### QUESTIONS?

#### Chronic Pain

- Affects approximately 20% of U.S. adults (50 million)
- Can limit an individual's ability to perform daily activities both in their personal and their professional lives
- Can lead to anxiety and depression
- Is difficult to treat successfully

#### Tolerance and the Opioid Epidemic

- People who suffer from chronic pain get opioids
- After a while, people become tolerant to the opioids
- Can lead to overdoses

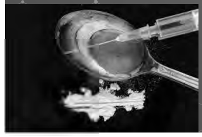

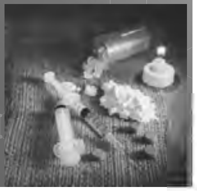

#### Potential Conclusions

Over time, alcohol tolerance should increase in flies, meaning the flies should see less negative effects at the same dose.

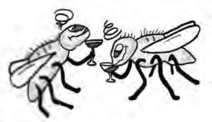

#### Pain

"A fifth vital sign"

"biopsychosocial phenomenon that includes sensory, emotional, cognitive, developmental, behavioral, spiritual, and cultural components"

Acute:

- Warns us of tissue damage
- Will heal and go away within 6 months

Chronic:

- Lasts longer than 6 months
- Can continue after the body has healed

#### Hypothesis

The pain and tolerance assays are both keen experiments that can help us better understand the physiological effects alcohol, opioids, and other addictive drugs have on the human body. This is because, in each experiment, the conductor must evaluate the ethanol related behaviors of the flies and see how ethanol can chemically and physically affect them. In the assays, the speed of how fast the flies can climb to the top of their sealed vial, after being exposed to ethanol, can help demonstrate how the flies react to the startling change in environment. However, prior to to conducting the experiment, a theory is that once the flies are exposed to the ethanol and are startled, their speed, while climbing to the top of the vial, will decrease after each trial because they will have gotten used to the effects of the ethanol.

#### Negative Geotaxis Assay

- Baseline assay to record the *Drosophila's* tendency to ascend upwards on the walls
- This behavior is an escape response for the *Drosophila*
- The escape response decreases as the flies age
- External factors do not alter results(density of flies being tested, the time of day, or repeated testing)

#### Data Collected

Painless (Vial #27895)
0.55621 (55.621%)

- Average of the flies crossing the 5 cm line

Wild Type (Vial #64349)
0.57725 (57.725%)

- Average of the flies crossing the 5 cm line

P-Value:
0.80679

#### *Drosophila Melanogaster*

- Also known as "fruit fly"
- Experiences pain & addiction
- Our experiments uses:
    - Wild Type (#64349)
    - Painless (#27895)

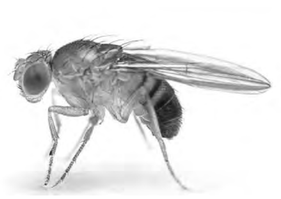

#### Any questions so far?

#### Hypotheses Created By Group 1

- Null hypothesis: Wild type negative geotaxis responses = Painless responses (i.e. there is no significant difference)
- Alternative: Wild type escape responses ≠ Painless escape responses

#### Opioids Today

| What we know: | What we don't know: |
|---|---|
| • In 2018, the average prescribing rate in the U.S. was 51.4 prescriptions per 100 people<br>• 21-29% of individuals prescribed opioids for chronic pain misuse them<br>• In 2018, 47,000 people died from opioid overdose<br> ○ 32% of these involved prescription opioids | • Why are people who suffer from chronic pain at greater risk to develop a substance use disorder?<br> ○ Are there biological, psychological and social aspects that increases the risk?<br> ○ Do the neurological circuits for chronic pain and reward overlap? |

#### How this research relates to this course

The two assays that will be further explained throughout this presentation go hand-in-hand with this course: "Neurology of Opioids and Pain". Due to the fact that this is a physiological based course, the pain and tolerance assays help further our understanding of how specimen, or in this case fruit flies, react to addictive drugs, such as opioids. However, in the case of this experiment, the role of opioids is played by ethanol. The research gathered by the two assays will help the conductor, as well as other soon-to-be scientists, gain more knowledge and theories about how addictive drugs and alcohol can affect a human, both mentally and physically, be observing the test fruit flies' ethanol-induced behaviors.

#### Negative Geotaxis Experimental Design

1. Carefully anesthetize one vial of flies on an ice pack
2. Transfer flies into a tube with a line 5 cm above the bottom
3. After 1 minute of flies adapting to environment, knock flies down to bottom by knocking vial onto a hard surface (3 hits)
4. Then put vial upright on a flat surface and start an 18 second timer
5. Count the amount of flies passing the line in those 18 seconds and repeat three trials for each vial of flies
6. Record data into the Data Presentation Excel Sheet

#### Sensitivity Assay

- Behavioral changes in mammals and flies due to alcohol being introduced to the system
- Genes have been identified in flies that influence physiological responses to alcohol
- Almost the same as the Negative Geotaxis Assay but with the inclusion of ethanol to the experiment with their escape response
- This assay will determine and better understand how the responses of the flies are affected by ethanol

#### Data Collected

Painless (Vial 27895) Averages

- 0%- 0.6968 (69.68%)
- 25%- 0.5782 (57.82%)
- 50%- 0.3779 (37.79%)

Wild Type (Vial 64349) Averages

- 0%- 0.8177 (81.77%)
- 25%- 0.4921 (49.21%)
- 50%- 0.3910 (39.10%)

Ttest Values:

- 0%- 0.2074738689
- 25%- 0.2952924
- 50%- 0.9134784558

#### Tolerance Assay

- Tolerance is defined as, "A condition that occurs when the body gets used to a medicine so that either more medicine is needed or different medicine is needed." by the National Cancer Institute.
- Examining the rapid-ethanol tolerance of the two genotypes of Drosophila will aid in further understanding how chronic pain is correlated to heightened addiction.
- This assay allows us to assess how the sensitivity of the Drosophila becomes affected when re-exposed to ethanol.
- This will allow us to gather information about the the tolerance that each genotype develops to the ethanol after a second exposure to both 0% and 50% ethanol.

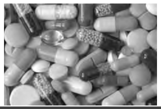

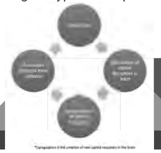

#### Group 1 Graph of Negative Geotaxis Assay

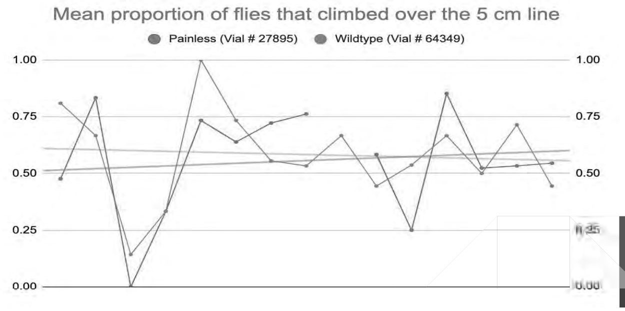

#### Hypotheses

Wild type (Vial 64349)

**Null hypothesis:** As alcohol concentration increases then so does the number of flies crossing the 5 cm line

**Alternative:** As alcohol concentration increases then there is a decrease in number of flies crossing the 5 cm line

Painless (Vial 27895)

**Null hypothesis:** Every group of painless flies do not cross the 5 cm line

**Alternative:** As alcohol concentration increases the amount of flies crossing the 5 cm line also increases

#### Graph Data for Sensitivity Assay

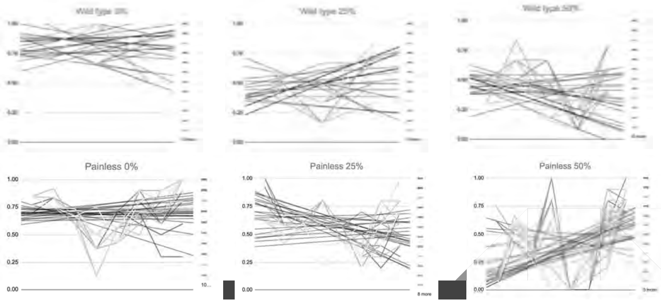

#### Conclusions

- Group 1 assumed a p-value less than 0.05 which any data collected below that would suggest it is statistically significant
- The p-value obtained of 0.80679 is not significant and fail to reject the null hypothesis
- The p-value demonstrates that the negative geotaxis should not be affected by pain

#### Sensitivity Assay Experimental Design

1. Anesthetize the flies in both vials and divide into three different groups for each vial (if excessive amounts of flies available)
2. Transfer one group of one vial into a tube with a 5 cm line marker
3. Cap the tube with a cotton plug doused in 0% (percentage of alcohol) hand sanitizer
4. Start 18 second timer and count how many flies pass the 5 cm line
5. Repeat process for 20 minutes with 18 sec timer and 1 minute rests
6. Then switch cotton plugs to 25% and 50% alcohol and repeat the same process with other groups who have not done the 0% cap
7. Record the 120 trials

#### Conclusions

- The ttest values of 0% and 25% show almost the two groups of flies are acting differently as the ttest values are very low
- 50% concentration group had a very high ttest value and almost acting the same in both groups
- The null hypothesis for both vials of flies can be failed to reject
- Ttest values show weirdly enough that 0% and 25% concentration, the flies act very differently but with the 50% almost act identical

#### Graphs for Tolerance Assay

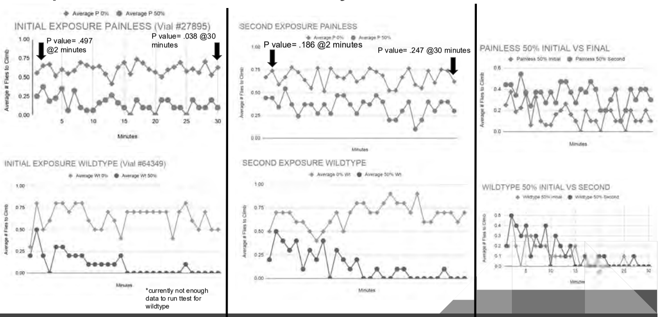

#### Experimental Design

1. Prepare 4 microcapillary pipettes by filling two with 0% ethanol, and two with 15% ethanol and two cotton plugs
2. Mark the starting volumes of each of the 4 pipettes using a marker, and make sure you can discern between the 0% and 15% ethanol pipettes for each genotype vial by either using a piece of tape or a marker.
3. Place one pipet with 0% and one with 15% into each of the cotton plugs
4. Anesthetize your flies and move all of the wild type flies into one empty vial, and all of the painless flies into a seperate empty vial. Then, use the two cotton plugs containing the pipettes to close the vials.
5. As soon as all the flies wake up, set a timer for 24 hours and begin to record data.
    a. Every hour, make a mark on the pipette so that after 24 hours there are 25 marks. This will allow you to calculate the volume and collect data following the 24 hour period.

#### Questions?

#### Hypotheses

Wildtype:

Hypotheses= The Drosophila will experience a decrease in sensitivity following the second exposure to ethanol.

Null Hypotheses= The Drosophila will experience heightened sensitivity following the second exposure to ethanol.

Painless:

Hypotheses= The Drosophila will experience experience no change in sensitivity following the second exposure.

Null Hypotheses= The Drosophila will experience an increase in sensitivity following the second exposure.

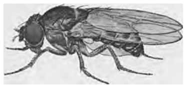

#### Cafe Assay

- This assay is used to demonstrate the addictive behaviors that Drosophila portray.
- It is known that the flies will self administer ethanol to pharmacologically relevant concentrations, therefore this assay will allow us to analyze addictive behaviors across both genotypes.
- Drosophila have tendencies to return to ethanol consumption after periods of abstinence which shows their addictive behavior.

#### Cafe Assay Predictions

- The wildtype flies will consume more ethanol than the painless flies because feeling pain makes them more susceptible to addiction
- The painless flies will consume an equal amount of 15% ethanol and 0% ethanol because it would have less of an effect on them.

#### Experimental Design

This experimental design is very similar to the design for the sensitivity assay. There are several differences.

1) In the tolerance assay, we used only the eppendorf tubes containing 0% and 50% ethanol
2) We recorded 30 trials instead of 20
3) After four hours, we repeated each of the steps and recorded data until the drosophila were no longer moving

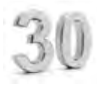

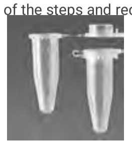

#### Conclusions

- At this moment, there are no definitive conclusions that can be made for the wild type flies
- The painless flies developed a tolerance as shown by the p values when comparing the initial and second exposure to 0% and 50% ethanol
    - Initial exposure p values over time indicate that the painless flies exposed to 50% ethanol had increased sensitivity
    - Second exposure p values over time indicate that the painless flies exposed to 50% ethanol acted similar to the flies exposed to 0% ethanol indicating that a tolerance was built up.
    - Our hypothesis was incorrect because the painless flies did experience a change in sensitivity following the second exposure
- Once we have more data for the wild type flies, we can analyze which genotype more rapidly developed a tolerance to the ethanol after being exposed to 50% ethanol twice

#### Hypotheses

Wildtype:

Hypotheses= Wild type flies will consume a greater amount of 15% ethanol than 0% ethanol.

Null Hypotheses= Wild type flies will consume the same amount of 15% and 0% ethanol.

Painless:

Hypotheses= Painless flies will consume the same amount of 15% ethanol and 0% ethanol.

Null Hypotheses= Painless flies will consume a greater amount of 15% ethanol than 0% ethanol.

### Thesis Presentation

#### Why do people rely on opioids and alcohol?

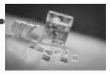

1. Most people rely on opioids to treat their chronic pain but roughly 21 to 29% of people prescribed opioids for chronic pain misuse them.*
2. About 80% of people who use heroin first misused prescription opioids. *
3. The presence of anxiety, depression, bipolar disorder, and other mental health issues increase the risk of alcoholism**
4. Drinking alcohol while on medication can lead to people to become addicted to the effects mixing substances has**

*National Institute on Drug Abuse. (2020, June 10). Opioid Overdose Crisis. Retrieved December 09, 2020, from https://www.drugabuse.gov/drug-topics/opioids/opioid-overdose-crisis

** 5 Most Common Causes of Alcoholism (And What to Do Next). (n.d.). Retrieved December 09, 2020, from https://talbottcampus.com/addiction-and-mental-health-resources/alcoholism-and-drug-abuse-articles/common-causes-alcoholism/

#### Protocol 1: Negative Geotaxis Assay

**Procedure:**

1. Two vials were used; one with the wild type flies and one with painless flies
2. The wild type flies were first anesthetized using an ice pack
3. The flies were then transferred to a new vial without food where they regained consciousness
4. After 1 minute, the vial was knocked on the table 3 times to induce the negative geotaxis response in flies
5. A timer was used to time for 18 seconds to see how many flies would cross the marking on the vial
6. Steps 25 were repeated 3 times for each group of flies

**Rationale:**

- "Negative geotaxis, an innate escape response during which flies ascend the wall of a cylinder after being tapped to its bottom, is one of the behaviors that senesces in *Drosophila*"
- We were able to see if there was a difference in the number of wild type and painless flies who climb over the line which would represent that a negative geotaxis response was elicited in the flies. helped to get a baseline understanding of how many flies demonstrate the negative geotaxis response without being exposed to anything

#### Sources continued

**Chronic pain**

- https://www.cdc.gov/mmwr/volumes/67/wr/mm6736a2.htm
- https://www.ncbi.nlm.nih.gov/pmc/articles/PMC4781803/
- https://www.drugabuse.gov/drug-topics/opioids/opioid-overdose-crisis
- https://www.sciencedirect.com/science/article/pii/S0896627315010338

#### Reasoning behind our study

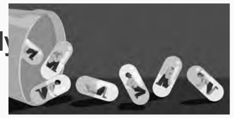

##### Why are we studying this?

In class, we discussed the effects that chronic pain can have on a person. Whether it be from an injury or a preexisting condition, chronic pain can lead to a person to be prescribed opioids as a way to treat this pain and discomfort they're feeling. One tragic effect of this opioid treatment is that most people are overprescribed opioids and end up being addicted and dependent on opioids. This addiction tends to snowball and lead to chronic pain patients trying other substances to feed their addiction and help their pain.

Does chronic pain increase the likelihood of someone becoming addicted to opioids and alcohol?

### Experimental Design and Results

#### Sources

- Proposal Presentation
- Group 1 Data Presentation 1
- Negative Geotaxis Assay
  - https://www.sciencedirect.com/science/article/abs/pii/S0531556505000343?via%3Dihub
- Sensitivity Assay
  - https://pubmed.ncbi.nlm.nih.gov/19645731/
- Tolerance Assay
- CAFE Assay
  - https://www.ncbi.nlm.nih.gov/pmc/articles/PMC2805771/

### Re Introduction

Why are we studying this?

What was our original hypothesis?

What protocols did we follow?

01.
##### Negative Geotaxis Assay
The purpose of this assay is to set the baseline of the escape response (climbing) for the flies before they're exposed to any substance.

02.
##### Sensitivity Assay
Learning how sensitive and tolerant the flies are to ethanol allows for us to see what genes are involved in ethanol related behaviors in the flies.

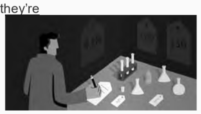

03.
##### CAFE Assay
Tracking how much sugar water and ethanol the flies drink helps establish a preference and helps show which one the flies prefer.

04.
##### Leg Amputation
This assay shows if amputation has any effect on the pre established preference for ethanol in the flies.

#### Negative Geotaxis Assay Data

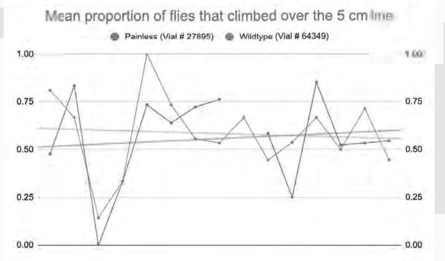

#### Sensitivity Assay Data (Wild Type)

| 0% Concentration | 25% Concentration | 50% Concentration |
|---|---|---|
| Trial 1- 0.76678 (76.678%)<br>Trial 10- 0.8 (80%)<br>Trial 20- 0.85 (85%) | Trial 1- 0.44433 (44.433%)<br>Trial 10- 0.49989 (49.989%)<br>Trial 20- 0.49433 (49.433%) | Trial 1- 0.49007 (49.007%)<br>Trial 10- 0.41333 (41.333%)<br>Trial 20- 0.29673 (29.673%) |

#### Protocol 2: Sensitivity/Tolerance Assay

**Procedure for Tolerance Assay:**

1. Starting with one genotype, the negative geotaxis experiment was done again but with 0% ethanol
2. 5-10 flies of the first genotype were used and after a minute rest period, the experiment was repeated 30 times
3. A four hour rest period was timed
4. During the four hour rest period 5-10 different flies of the same genotype were put through the negative geotaxis experiment but with 50% ethanol this time.
5. After the four hour rest period, the 0% and 50% cotton plugs were resaturated and the sensitivity assay was observed again
6. These steps were repeated for the second genotype

**Rationale:**

- Since fruit flies have acute sensitivity to ethanol they also have a rapid ethanol tolerance
- This experiment allowed us to see how the flies were affected after being re-exposed to ethanol and developing a tolerance to the ethanol and if that had an effect on their negative geotaxis response
- This experiment also allowed us to see how the negative geotaxis response is affected due to tolerance in wild type and mutated painless flies.

#### Negative Geotaxis Assay Data

| Wild type (Vial #64349) | Painless (Vial #27895) |
|---|---|
| 0.57725 (57.725%) | 0.55621 (55.621%) |
| Mean of flies crossing the 5 cm line | Mean of flies crossing the 5 cm line |

| Difference of Means | p-value |
|---|---|
| ±0.02104 | 0.80679 |

#### Sensitivity Assay Hypotheses

##### Wild Type

**Null Hypothesis:**
As alcohol concentration increases, the number of flies crossing the 5 cm line increases.

**Alternative:**
As alcohol concentration increases, the number of flies crossing the 5 cm line decreases.

##### Painless

**Null Hypothesis:**
Every group of painless flies does not cross the 5 cm line.

**Alternative:**
As alcohol concentration increases, the number of flies crossing the 5 cm line increases.

#### Sensitivity Assay Data

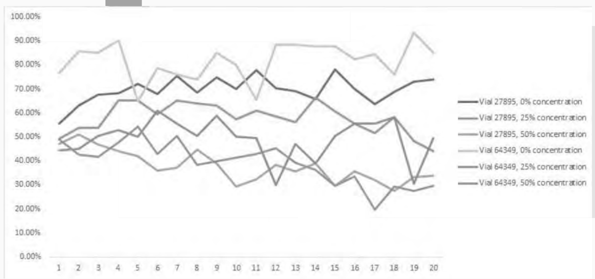

#### Negative Geotaxis Assay Hypotheses

**Null Hypothesis:**
Wild type response = Painless response
(i.e. there is no significant difference)

**Alternative:**
Wild type escape responses ≠ Painless escape responses

#### Protocol 2: Sensitivity/Tolerance Assay

Procedure for Sensitivity Assay:

1. Both groups of flies were anesthetized on an ice pack
2. The flies were again transferred to new vials marked with 5cm lines
3. The new vials were capped with a cotton plug 0% alcohol (hand sanitizer)
4. The vials were knocked against the table and the times was started for 18 seconds to see how many flies climbed pass the line in 18 seconds
5. This was repeated 20 times
6. Step 4 was repeated with a 25% and 50% cotton plug for a total of 20 trials for each percentage of alcohol exposure for both of two groups of flies

**Rationale:**

- **Fruit flies have an intrinsic ability of sensing alcohol and its** easy to see the effect of alcohol on fruit flies because they display simple alcohol-induced behaviors such as sedation and motor impairment.
- This experiment was done to see if alcohol exposures and different percentages of alcohol causes an increased negative geotaxis response in flies causing them to be more drawn to the top of the vial because they sense the alcohol.
- "Fruit flies have a number of genes have been identified that influence physiological responses to ethanol."
- Wanted to test the acute sensitivity to ethanol that fruit flies have

#### Sensitivity Assay Data (Painless)

| 0% Concentration | 25% Concentration | 50% Concentration |
|---|---|---|
| Trial 1- 0.55479 (55.479%)<br>Trial 10- 0.69904 (69.904%)<br>Trial 20- 0.74038 (74.038%) | Trial 1- 0.49191 (49.191%)<br>Trial 10- 0.57266 (57.266%)<br>Trial 20- 0.442 (44.2%) | Trial 1- 0.47183 (47.183%)<br>Trial 10- 0.29379 (29.379%)<br>Trial 20- 0.33963 (33.963%) |

### Protocol 3: CAFE Assay

**Procedure:**
1. Four microcapillary pipettes were prepared; two with 0% ethanol and two with 15% and two cotton plugs with nothing on them
2. The starting volumes of each of the pipettes was marked
3. One 0% and one 15% pipette were placed into each of the cotton plugs
4. The flies were anesthetized and the two different genotypes were placed in different vials and were capped with the two cotton plugs containing the pipettes
5. Once the flies regained consciousness, a timer for 24 hours was started
6. Each hour it was observed how much volume was remaining in the microcapillary pipettes

**Rationale:**
- This experiment helped to see the addictive behaviors of fruit flies
- "Flies prefer to consume ethanol-containing food over regular food, and this preference increases over time. Flies are attracted to the smell of ethanol, and start to exhibit features of alcohol addiction."
- "First, flies self-administer ethanol to pharmacologically relevant concentrations. Second, flies overcome an aversive stimulus in order to consume ethanol. Third, flies rapidly return to high levels of ethanol consumption after a period of imposed abstinence."

### Protocol 4: Leg Amputation

**Procedure:**
1. A middle appendage amputation was performed on the fruit flies just below the femur
2. Flies were recovered for one day prior to performing the negative geotaxis assay

**Rationale:**
- Nociception is the sensory process by which animals can detect and avoid harmful **stimuli**
- **Neuropathic allodynia is pain due to a stimulus that does not normally produce pain**
- Nerve injury such as an amputation "triggers a loss of central inhibition that drives escape circuit plasticity and neuropathic allodynia."
- From this experiment we were able to see if chronic pain caused by the amputation leads to ethanol sensitivity.

### Leg Amputation: Negative Geotaxis Assay Data

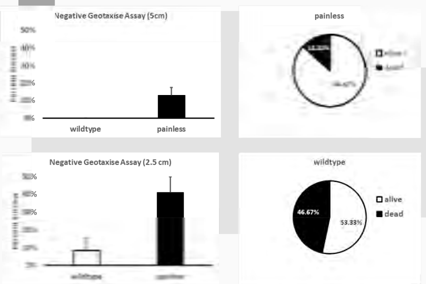

### Tolerance Assay Data

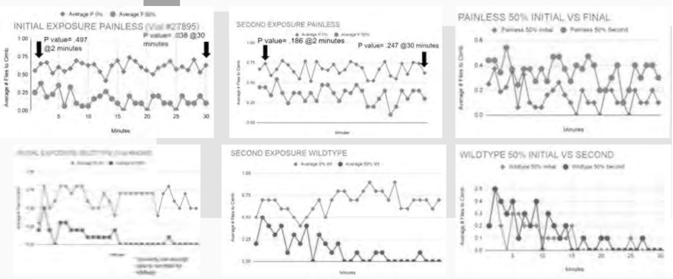

### CAFE Assay Data

No conclusions can be drawn since very little data was collected :(

### Leg Amputation: Negative Geotaxis Assay Results

- Painless flies have increased mobility after amputation.
- 5 cm is too much for all flies after amputation.
- Painless flies survive at a greater rate after amputation.

### Tolerance Assay Hypotheses

#### Wild Type

**Hypothesis:**
The flies will experience a decrease in sensitivity after the second exposure to ethanol.

**Null Hypothesis:**
The flies will experience heightened sensitivity after the second exposure to ethanol.

#### Painless

**Hypothesis:**
The flies will experience no change in sensitivity after the second exposure to ethanol.

**Null Hypothesis:**
The flies will experience an increase in sensitivity after the second exposure to ethanol.

### CAFE Assay Hypotheses

#### Wild Type

**Hypothesis:**
Wild type flies will consume a greater amount of 15% ethanol than 0% ethanol.

**Null Hypothesis:**
Wild type flies will consume the same amount of 15% and 0% ethanol.

#### Painless

**Hypothesis:**
Painless flies will consume the same amount of 15% and 0% ethanol.

**Null Hypothesis:**
Painlessflies will consume a greater amount of 15% ethanol than 0% ethanol.

### Leg Amputation: Negative Geotaxis Assay

**Hypothesis:**
Amputation should not affect mobility differently.

**Prediction:**
Flies in each group will climb at similar rates.

#### Leg Amputation: Sensitivity Assay Data

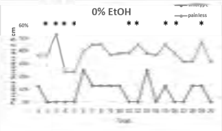

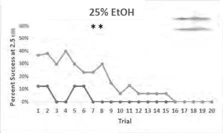

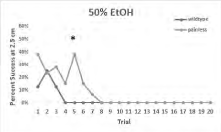

#### Where can we go with this in the future?

-Repeat assays, to see if consistency remained the same or what differed

- Test other substances like opioids, to see if there is a difference in results

-We could assess whether the increased tolerance is dose dependent

-To possibly increase the validity of the leg amputation assay, we could somehow design a way for the flies to have impaired coordination

-Design control experiment so flies have impaired coordination and compare results

-Create a sensation of pain that does not result from amputation, and that does not affect mobility

-We could more forward with developing future experiments that show other direct connections between chronic pain the increased likelihood of addiction

#### Leg Amputation: Sensitivity Assay Results

- Painless flies were less sensitive to 25% and 50% ethanol.

#### Conclusions

Negative Geotaxis Assay
- Can't reject null hypothesis

Sensitivity Assay
- Low t test values: 0%, 25%
- High t test values: 50%

Tolerance Assay
- Painless flies had increased sensitivity at initial exposure,
- Tolerance at second exposure

Leg Amputation Assay
- Painless flies had better mobility, despite hypothesis
- Seen at the 5cm mark
- Persisted at 2.5 cm mark
- Conclusion was significant because this was not expected

#### Leg Amputation: Sensitivity Assay

Hypothesis:
Pain will increase sensitivity to ethanol.

Prediction:
Wild type flies will have increased sensitivity to ethanol.

### Conclusions

#### Questions?

### APPENDIX 10 – JC PRESENTATION

### Study On Nerve injury and Neuropathic Sensitization in Drosophila

### Introduction/Background

#### The Field

- The field that is being studied here how fruit flies that experience injury may develop chronic pain by damaging nociception causing allodynia
- Nociception has developed over the past 500 million years and helps animals avoid a situation that make cause harm, however damage to nociception can lead to an adverse effect of its intended purpose
- What is not known about the field is how and why damage to our nociception occurs on a molecular level.
- So this experiment will try and answer that very question.

#### Research Questions

- What are the molecular mechanisms responsible for neuropathic sensitization (hypersensitivity to innocuous stimuli that can cause chronic pain; i.e. allodynia) that develops as a result of nerve injury in *Drosophila melanogaster* (fruit flies)?
- In order to answer their main question, the researchers first had to know if nerve injury in fact leads to allodynia in *Drosophila*.

### Where Does It Fit?

#### The Basic Connection

- In class, we are covering topics that are directly related to the nervous system
  - Neuroanatomy
  - Neurophysiology
  - Pain Neurobiology
- Our course name is "Neuroscience in the Opioid Epidemic"
  - Lack of effective treatments for chronic pain has had knock-on effects in our society, specifically the opioid epidemic

#### A Focus On Chronic Pain

- Despite decades of research into the molecular and physiological mechanisms that contribute to neuropathic pain, it is still not completely clear what we should target to treat the underlying pathology
- Chronic pain in the fly is caused by the loss of sensory neurons in the leg, and is a common cause of neuropathic pain in humans, according to the study
- The loss of central inhibition spinal cord (VNC) leads to neuropathic allodynia and a change in the nociceptive escape circuit physiology, all of which are hallmarks of human pain

#### A Focus On Chronic Pain

- Together, the data highlight a previously unknown neuropathic injury response program that promotes heightened sensory vigilance and an augmented escape response changes that may help promote survival in dangerous environments.
- Similar to the observations in the experiment, painful peripheral neuropathy in response to the chemotherapy agent vinblastine also requires TrpA1 both in flies and mice, suggesting that the underlying mechanism of sensitization in response to different injuries may show some conservation.

#### Chronic Pain - Working With Patients

- Current therapies do not adequately address pain for most patients
- Neuropathic pain is generally refractory to available therapies, with fist-line antineuropathic providing adequate pain relief for only ~25% of patients
- We find that peripheral nerve injury leads to central disinhibition, neuropathic allodynia and nociceptive hypervigilance in the fly
  - Now with this information, this research helps us to take the next steps to working with human patients in our community
- A basic understanding of the conserved architexture driving neuropathic pain can help create better and non-addictive pain therapies to reverse or resolve chronic pain

#### Detecting Allodynia

- In determining if *Drosophila* developed allodynia after nerve injury, researchers observed the average number of escape responses ("jumps") that injured and uninjured flies displayed on a hot plate at a specific temperature in a given timeframe (see Figure A).
- If there were a significantly greater number of jumps by the injured flies compared to the uninjured flies at the same temperature, it would indicate they are exhibiting pain responses to a stimuli normal *Drosophila* experience as sub-noxious, meaning injured flies have developed thermal allodynia.

### Results

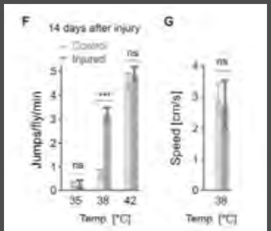

#### Injury Model

- Following control tests, flies were subjected to bodily injury; amputation below the femur of the middle-right leg
  - See Figure C:

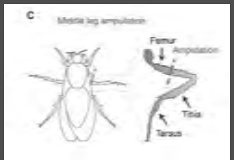

#### Investigation into Causes

- Researchers examined changes in the nociception circuitry of nerve-injured *Drosophila* using recordings of electrophysiological responses as well as imaging of the flies' nervous system to examine how neural functioning and pain circuitry change as a result of injury leading to allodynia (see figures 4F and 6D for examples).
- In order to determine which molecular mechanisms were behind the changes, researchers manipulated or interfered with specific neurological functions and genetic expressions to determine their roles in the development of allodynia following nerve injury by observing how the manipulation affected the development of allodynia.

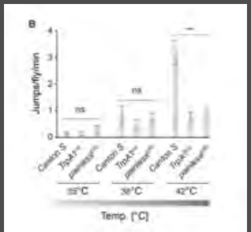

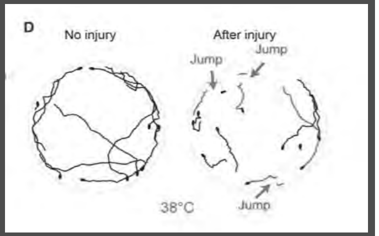

### Methods

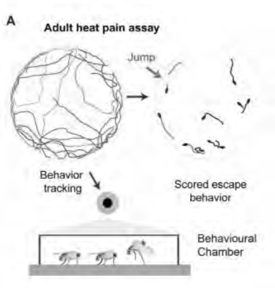

#### Initial Findings

- Flies showed significant escape responses at temperatures greater than 42 degrees Celsius (See Figure B).
- Following amputation, injured flies showed significant higher escape response at temperatures approaching 38 degrees (See Figures D and F).
  - Responses to the critical temperature of 42 degrees celsius did not alter significantly
    - Injured flies also showed a slight increase in the speed of their escape attempts (See Figure G).

#### Further Experimentation

- Following the initial tests, researchers attempted to test fly Allodynia after injury.
  - Researchers used *UAS-tetanus toxin* to prevent the response of the *ppk+* sensory neurons
    - The blockage of response prevented thermal Allodynia from forming.
  - Researchers also interfered with *TrPA1* expression, which prevented Allodynia as well.
    - The researchers concluded that the expression of *TrPA1* in the *ppk+* sensory neuron was required for Allodynia

#### GABA's Role

- The researchers then examined how the reduction in GABA lead to neuropathic pain.
  - After injury, flies lost 40% of GABA activity in the injured area.
    - Scientists attempted to reduce the loss by blocking the role of caspase, a molecule used in the process of cell death
    - Following the use of caspase, sensitivity to thermal stimulation was completely blocked
      - The prevention of apoptosis in GABA cells prevented the formation of neuropathic allodynia

#### Figure 4f and 4h

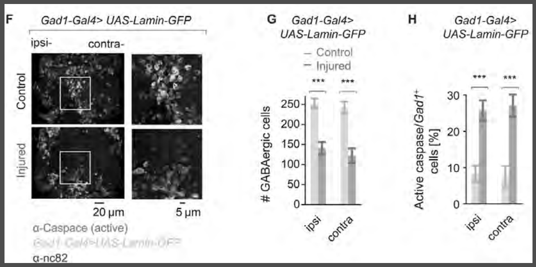

#### Twist protein Figures 6D and 6F

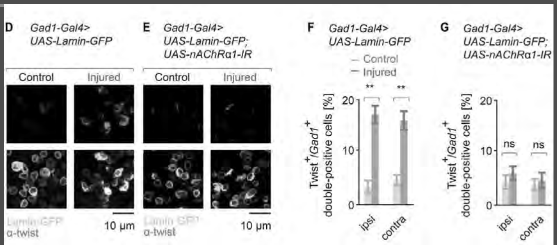

### Conclusions

#### Conclusion

- The results of the study show that when an organism is in chronic pain, they will have a hyper sensitivity to pain and will be more alert to danger
- The results support the hypothesis because they neuropathic pain can lead to an animal being more vigilant and sensitive to allodynia for survival
- The outcome of the study show that molecular response is the cause of the allodynia in insects

### Questions/Discussions?

#### Questions

1. Could similar methods such as the use of chemicals to prevent cell death in an injured area be used as a non-addictive chronic pain relief mechanism for humans?
2. What are potential studies that could be used in order to test human individuals with pre-existing nerve injury with their vigilance and neuropathic sensitization?

#### APPENDIX 11 – CONFERENCE DEBRIEF PRESENTATION

#### Introduction: Field

- "Pro-bdnf Changes in the Fronto-cerebellar Circuitry after Oxycodone -induced place Preference in Adult Rats"
- Research conducted by Alejandro Torres, Zinar Simsek and Dolores Vazquez Sanroman, PhD, of Oklahoma State University, Center for Health Sciences
- Neuroscience, sociology, public or population health, public policy, mental health
- Cerebellum: motor functions

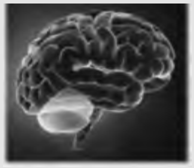

### How does this fit in?

### Experimental Approach

### Introduction

#### Introduction: Researcher's Questions

- How does oxycodone preference affect the cerebellum and prefrontal cortex?
- About oxycodone
  - Brand names: Dazidox, Eth-Oxydose, Oxaydo, OxyCONTIN, OxyCONTIN CR, Oxydose, Oxyfast, Oxy IR, Roxicodone, and Roxicodone Intensol.
- About prefrontal cortex
  - El-Baba and Schury (2020): "The prefrontal cortex is known to be the higher-order association center of the brain as it is responsible for decision making, reasoning, personality expression, maintaining social appropriateness, and other complex cognitive behaviors."
- About BDNF
  - Medline: "The BDNF protein is found in regions of the brain that control eating, drinking, and body weight; the protein likely contributes to the management of these functions."
- Implications for substance use disorder and biological mechanisms of addiction

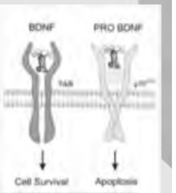

#### Methods

##### Conditioned Place Preference (CPP)

PND120, male adult rats were tested for CPP after being randomly assigned to the oxycodone drug group or the vehicle group. Following conditioning, rats were given a 20 min CPP test evaluating oxycodone preference.

##### Tissue Homogenization

Tissue was placed in microcentrifuge tube with 800ul of RIPA buffer and lysed using the ultrasonicator. Centrifuged for 20 min at 14,000rpm before transferring the supernatant into a new tube and kept in the freezer at -80°C.

##### Brain Extraction/Dissection

Rats were euthanized using the $CO_2$ chamber. Brains were carefully removed and quickly placed in $LN_2$ and kept at -80°C before sectioning. The cerebellum was dissected from the brain in vermis sections.

##### BCA Protein Assay and ELISA

Tissue lysates were diluted using RIPA buffer; BCA Protein Assay Kit was used. For immunoassay, 96 well plates and biosensis mature and proBDNF Rapid ELISA kit was used to analyze diluted cerebellar lysates in duplicates. OD was read at 450nm.

### P951- Pro-bdnf Changes in the Fronto-cerebellar Circuitry after Oxycodone-induced place Preference in Adult Rats

#### Introduction: Unknowns and Developments in Field

- Koziol et al. (2014): more research needed to confirm role of cerebellum in cognitive functioning; appears to have a role in "neurocognitive development, language function, working memory, executive function, and the development of cerebellar internal control models," but these are inferences.
- Cognitive functioning and substance use disorders
- Example of research put into practice: Medication Assisted Treatment
  - Substance Abuse and Mental Health Services Administration: MAT "operates to normalize brain chemistry, block the euphoric effects of alcohol and opioids, relieve physiological cravings, and normalize body functions without the negative and euphoric effects of the substance used." Buprenorphine, methadone, and naltrexone are approved for opiate use disorder.
  - Behavioral Health Group: buprenorphine, methadone are opioid agonists; naltrexone is an opioid antagonist

- There is a lot still unknown about understanding opioid addiction
- This research helps to further understand:
  - mechanisms behind opioid addiction, in this case the effects of oxycodone
  - how the brain responds to opioids → pro-BDNF levels
  - how the brain restores itself after opioid use
  - how different parts of the brain are affected→ fronto-cerebellar circuitry

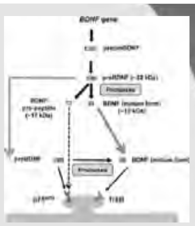

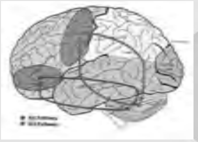

#### Main Result:

##### Oxycodone creates a conditioned place preference in male adult rats.

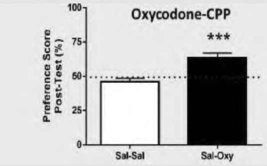

- Oxycodone-treated animals acquired CPP : The animals spend more time in the drug paired chamber which indicates pre-anticipation of stimulus.

**Fig1.** Oxycodone induced CPP [ F (1, 21) = 55.02, p < .001]

### Conclusion

#### References

Behavioral Health Group. Medication assisted treatment. (MAT). https://bhgrecovery.com/treatment-services/proven-treatment/.

Behavioral Health Group. About us. https://bhgrecovery.com/about-us/.

Medline Plus. BDNF gene. NIH. https://medlineplus.gov/genetics/gene/bdnf/.

Medline Plus. Cerebellum - function. NIH. Reviewers: Minagar, A and Zieve, D. https://medlineplus.gov/ency/imagepages/18008.htm.

El-Baba RM, Schury MP. Neuroanatomy, Frontal Cortex. [Updated 2020 Jul 31]. In: StatPearls [Internet]. Treasure Island (FL): StatPearls Publishing; 2020 Jan-. Available from: https://www.ncbi.nlm.nih.gov/books/NBK554483/.

Elmore S. (2007). Apoptosis: a review of programmed cell death. Toxicologic pathology, 35(4), 495–516. https://doi.org/10.1080/01926230701320337.

Koziol, L. F., Budding, D., Andreasen, N., D'Arrigo, S., Bulgheroni, S., Imamizu, H., Ito, M., Manto, M., Marvel, C., Parker, K., Pezzulo, G., Ramnani, N., Riva, D., Schmahmann, J., Vandervert, L., & Yamazaki, T. (2014). Consensus paper: the cerebellum's role in movement and cognition. Cerebellum (London, England), 13(1), 151–177. https://doi.org/10.1007/s12311-013-0511-x.

Mayo Clinic (2020). Oxycodone (oral route). https://www.mayoclinic.org/drugs-supplements/oxycodone-oral-route/description/drg-20074193.

OSU Center for Health Sciences. https://medicine.okstate.edu/.

Substance Abuse and Mental Health Services Administration. Medication assisted treatment (MAT).

### Results

#### Oxycodone induced CPP levels had decreased levels of pro-BDNF on the cerebellum

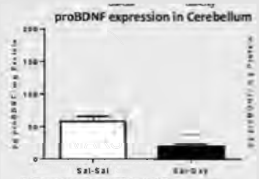

**Fig2.** Oxycodone decreased levels of proBDNF in the cerebellum [illegible]

The graph shows a decrease in the levels of pro-BDNF in the cerebellum of rats with an oxycodone CPP

- The protein expression of proBDNF in the cerebellum significantly decreased in rodents after oxycodone-induced CPP, while in the prefrontal cortex the levels of proBDNF significantly increased in comparison to saline control group.

#### Where can we go with this in the future?

- Investigate the fronto-cerebellar relationship between the levels of mature and proBDNF from other regions of the brain involved in reward seeking behavior
- Investigate whether adverse childhood experiences influence the onset of oxycodone addiction
  - Bio-psychesocial aspect of pain and addiction

- To mimic oxycodone induced addiction, an in vivo model consisting of adult Sprague Dawley male rats and the CPP paradigm was used.
- The rats received alternating injections of oxycodone and saline to pair an environment with the rewarding stimulus followed by an evaluation of CPP for oxycodone-seeking behavior.
- Lastly, the brain is extracted and dissected to analyze for mature and pro-BDNF levels.

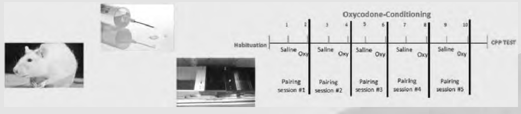

#### Oxycodone induced CPP increased levels of pro-BDNF in the prefrontal cortex

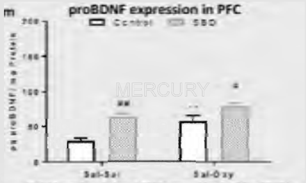

**Fig3.** Oxycodone induced CPP increased levels of Pro-BDNF in the PFC [illegible]

The graph shows a higher percentage of pro-BDNF in the prefrontal cortex of the rats with a oxycodone CPP

The increase in pro-BDNF in the prefrontal cortex helps focus the brain on the oxycodone rewarding properties that BDNF creates

- Oxycodone-induced conditioned place preference (CPP) modified Pro-BDNF levels in a "seesaw" mechanism in the fronto-cerebellar circuitry.
- A seesaw mechanism is one in which the brain restores neurotransmitters after opioid exposure; e.x. Alcohol exposure

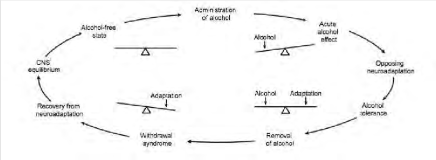

- Thus, pro-BDNF may be a required neurotrophin in the opioid-induced reorganization of fronto-cerebellar circuitry and fronto-cerebellar dysfunction.

### Questions?

1. What are some biopsychosocial factors that can influence oxycodone addiction?

1. Why might studying more about how addiction affects the cerebellum help our understanding of addiction? How could this lead to new, more effective treatments?

### APPENDIX 12 – SURVEY INSTRUMENT DESIGN & DETAILS

**Experimental Design:** We built a pre-/post-test design that utilized two validated instruments. All survey instruments were approved by the Rutgers Camden Institutional Review Board and provided informed consent for all participants (Rutgers ID: Pro2020002151). The first instrument, published by Stets et al., uses 11 questions with a 5-7 likert scale. Question 1 is used as a measure of science identity. The average response to Questions 2-7 measure Science Identity Prominence which is defined as how much the student values the importance of their science identity. The average of Questions 8-11 measure their reflected appraisal which is defined as how the student believes other's see them as a science student. The Science Identity Discrepancy is measured by subtracting the student's Science Identity from their Reflected Appraisal (i.e., more positive values indicate the Reflected Appraisal is greater than their Science Identity). The second instrument, published by Weston et al., is the Undergraduate Research Student Self-Assessment (URSSA) which has been adopted by several entities to assess whether specific research-related activities benefit a student's research goals.

This URSSA uses a 5-point likert scale where the average ratings of questions 1-8 measure the category of "thinking and working like a scientist", questions 9-14 measure "personal scientific gains", questions 15-26 measure "improvement of scientific skills", and questions 27-34 measure "improvement of researcher attitudes/behaviors".

The pre-test time point was placed at week 4 of the semester to assess what student's perceived as their gains from the first four weeks of class prior to receiving their lab in a box. At this time point, students would have been exposed to the course content, literature, and the experimental plan but they would not have received the "lab in a box" yet. We found this to be a better baseline than the beginning of class so that we could differentiate between gains from the course material vs the research experience. The post-test time point was during week 15.

We also measured the student's self-reported confidence in research activities at the pre- timepoint to identify if there would be any confounding factors that may make research gains more likely for some students. We assessed this as the average response to a 5-point likert scale of 15 questions designed by Stet et al.

**Science Identity Questionnaire:**

Using a scale of 1-7 (far above average, moderately above average, slightly above average, average, slightly below, moderately below, far below) ***(Science Identity)***

1. As a science student, I rate myself:

Using a scale of 1-5 (strongly agree, somewhat agree, neither agree or disagree, somewhat disagree, strongly disagree) ***(Science Identity Prominence)***

2. In general, being a scientist is an important part of my self-image.
3. I have a strong sense of belonging to the community of scientists.
4. Being a scientist is an important reflection of who I am.
5. I have come to think of myself as a ˜scientist".
6. I intend to pursue a science-related research career.
7. I intent to pursue a science-related non-research career.

Using a scale of 1-7 (far above average, moderately above average, slightly above average, average, slightly below, moderately below, far below) ***(Science Identity Discrepancy)***

8. How do you think that your family members rate you as a student?
9. How do you think your coworkers rate you at as a student?
10. How do you think your friends rate you as a student?
11. How do you think your partner/spouse rates you as a student?

**URSSA Questionnaire:**

Using a scale of 1-5 (great gains, good gains, moderate gains, minimal gains, no gains at all, no answer)

**(thinking and working like a scientist)**

1. Analyzing data for patterns.
2. Figuring out the next step in a research project.
3. Problem-solving in general.
4. Formulating a research question that could be answered with data.
5. Identifying limitations of research methods and designs.
6. Understanding the theory and concepts guiding my research project.
7. Understanding the connections among scientific disciplines.
8. Understanding the relevance of research to my course work.

**(personal scientific gains)**

9. Confidence in my ability to contribute to science.
10. Comfort in working collaboratively with others.
11. Confidence in my ability to do well in future science courses.
12. Ability to work independently.
13. Developing patience with the slow pace of research.
14. Understanding what everyday research work is like.

**(improvement of scientific skills)**

15. Writing scientific reports or papers.
16. Making oral presentations.
17. Defending an argument when asked questions.
18. Explaining my project to people outside my field.
19. Preparing a scientific poster.
20. Keeping a detailed lab notebook.
21. Conducting observations in the lab or field.
22. Using statistics to analyze data.
23. Calibrating instruments needed for measurement.
24. Understanding journal articles.
25. Conducting database or Internet searches.
26. Managing my time.

**(improvement of researcher attitudes/behaviors)**

27. Engage in real-world science research.
28. Feel like a scientist.
29. Think creatively about the project.
30. Try out new ideas or procedures on your own.
31. Feel responsible for the project.
32. Work extra hours because you were excited about the research.
33. Interact with scientists from outside your school.
34. Feel a part of a scientific community.

**Research Confidence Questionnaire:**

Using a scale of 1-5 (strongly agree, somewhat agree, neither agree nor disagree, somewhat disagree, strongly disagree) ***(Research Confidence)***

1. I would feel really good if I were the only one who could answer the teachers' question in class.
2. It's important to me that the other students in my classes think that I am good at my work.
3. I want to do better than other students in my classes.
4. I would feel successful in school if I did better than most of the other students.
5. I would like to show my teachers that I'm smarter than the other students in my classes.
6. Doing better than other students in school is important to me.
7. It's very important to me that I don't look stupid in my classes.
8. An important reason I do my school work is so that I don't embarrass myself.
9. The reason I do my school work is so my teachers don't think I know less than others.
10. The reason I do my school work is so others won't think I'm dumb.
11. One reason I would not participate in class is to avoid looking stupid.
12. One of my main goals is to avoid looking like I can't do my work.
